# A screen of drug-like molecules identifies chemically diverse electron transport chain inhibitors in apicomplexan parasites

**DOI:** 10.1101/2022.02.13.480284

**Authors:** Jenni A. Hayward, F. Victor Makota, Daniela Cihalova, Esther Rajendran, Soraya M. Zwahlen, Laura Shuttleworth, Ursula Wiedemann, Christina Spry, Kevin J. Saliba, Alexander G. Maier, Giel G. van Dooren

**Affiliations:** Research School of Biology, Australian National University, Canberra, ACT, Australia

**Author notes:** J.A. Hayward and F.V. Makota contributed equally to this paper.

**Keywords:** electron transport chain, inhibitors, mitochondrion, Apicomplexa, *Plasmodium falciparum*, *Toxoplasma gondii*

## Abstract

With the advent of resistance to existing treatments, new drugs are needed to combat apicomplexan parasites such as the causative agents of malaria (*Plasmodium* species) and toxoplasmosis (*Toxoplasma gondii*). To identify new inhibitors of the mitochondrial electron transport chain (ETC) in these parasites, we developed a Seahorse XFe96 flux analyzer approach to screen compounds from the Medicines for Malaria Venture ‘Pathogen Box’ for ETC inhibition. We identified six chemically diverse, on-target inhibitors of the ETC of *T. gondii*, five of which also target the ETC of *Plasmodium falciparum*. Two of the identified compounds (MMV024937 and MMV688853) represent novel ETC inhibitor chemotypes. We pinpoint the molecular targets of these inhibitors, demonstrating that all target ETC Complex III, with MMV688853 additionally targeting a kinase with a key role in parasite invasion of host cells. Most of the compounds remain effective inhibitors of parasites that are resistant to the clinically used Complex III inhibitor atovaquone. In sum, we have developed a versatile screening approach to identify and characterize new inhibitors of the ETC in apicomplexan parasites.

## Introduction

Apicomplexan parasites cause numerous diseases in humans and livestock worldwide. Up to a third of the global human population is chronically infected with *Toxoplasma gondii*, which can cause the disease toxoplasmosis in immunocompromised or pregnant individuals (Montoya and Liesenfeld, 2004). *Plasmodium* parasites cause the disease malaria, which killed an estimated 409 000 people and infected 229 million in 2019 (WHO, 2020). Despite the recent approval of the first malaria vaccine for children by the World Health Organization, there is currently no effective vaccine against malaria for adults or against toxoplasmosis in humans. There is therefore a heavy reliance on drugs to treat both diseases. Current treatment options are limited and have questionable efficacy and safety. For instance, while frontline therapeutics such as pyrimethamine and sulfadiazine are able to kill the disease-causing tachyzoite stage of *T. gondii,* they fail to eradicate the long-lived bradyzoite cyst stage that causes chronic infection and elicit adverse effects in many patients (Alday and Doggett, 2017). Emerging resistance to frontline therapeutics, such as artemisinin, is a particular problem for treating the potentially life-threatening severe malaria caused by *Plasmodium falciparum* (Fairhurst and Dondorp, 2016). New treatments for toxoplasmosis and malaria are therefore much needed.

The mitochondrion is important for apicomplexan parasite survival and is a target of many anti-parasitic compounds (Goodman et al., 2017). Like in other eukaryotes, the inner membrane of the parasite mitochondrion houses an electron transport chain (ETC), which is composed of a series of protein complexes that contribute to energy generation and pyrimidine biosynthesis (Hayward and van Dooren, 2019). Electrons derived from parasite metabolism are fed into the ETC via the action of several dehydrogenases – including succinate dehydrogenase (SDH), malate-quinone oxidoreductase (MQO), glycerol 3-phosphate dehydrogenase (G3PDH), dihydroorotate dehydrogenase (DHODH), and type II NADH dehydrogenases (NDH2) – which all reduce the hydrophobic inner membrane electron transporting molecule coenzyme Q (CoQ). CoQ interacts with ETC Complex III (also known as the coenzyme Q:cytochrome *c* oxidoreductase or *bc*_1_ complex) at the so-called Q_o_ and Q_i_ sites, where electrons are donated to or accepted from Complex III, respectively, in a process termed the Q cycle (Mitchell, 1975). This process also contributes to the generation of a proton motive force across the inner mitochondrial membrane by transporting protons from the matrix into the intermembrane space. Complex III passes electrons to the soluble intermembrane space protein cytochrome *c* (CytC). CytC shuttles the electrons to ETC Complex IV (cytochrome *c* oxidase), which donates them to the terminal electron acceptor, oxygen. Complex IV also contributes to the proton motive force by translocating protons across the inner mitochondrial membrane. The net reaction of the ETC is thus the oxidation of cellular substrates and reduction of oxygen, coupled to the translocation of protons from the matrix into the intermembrane space to generate a proton gradient across the inner membrane. This proton gradient can be utilized by an F-type ATPase (Complex V) to generate ATP and for important mitochondrial processes such as protein import (Schmidt et al., 2010). In the erythrocytic stages of the *P. falciparum* lifecycle, the ETC functions primarily as an electron sink for the DHODH reaction in the *de novo* pyrimidine biosynthesis pathway rather than for ATP synthesis (Painter et al., 2007).

ETC Complex III is the target of many anti-parasitic agents, including the clinically used therapeutic atovaquone and the pre-clinical ‘endochin-like quinolone’ (ELQ) compounds (Fry and Pudney, 1992, Doggett et al., 2012, Stickles et al., 2015). Many Complex III-targeting compounds are CoQ analogs that bind to the Q_o_ and/or Q_i_ sites of Complex III (Barton et al., 2010). The ability of these compounds to selectively target parasite Complex III lies in differences in the CoQ binding site residues between parasites and the mammalian hosts they infect, specifically in the cytochrome *b* protein of the complex (Vaidya et al., 1993, Fisher et al., 2012, Fisher et al., 2020). For instance, the Q_o_ site inhibitor atovaquone has an IC_50_ value in the nanomolar range against Complex III activity in *T. gondii* and *P. falciparum*, but inhibits the mammalian complex 13- to 230-fold less effectively (Siregar et al., 2015, Nilsen et al., 2013, Doggett et al., 2012). Although it is a potent and selective inhibitor of Complex III in apicomplexans, resistance to atovaquone can readily emerge as the result of mutations in the cytochrome *b* protein (McFadden et al., 2000, Srivastava et al., 1999), limiting its use in treating the diseases caused by these parasites. Identifying Complex III inhibitors that remain effective against atovaquone-resistant parasites is therefore desirable.

Strategies to identify new anti-parasitic compounds often use high throughput screening of small molecule libraries to identify inhibitors of parasite proliferation (Smilkstein et al., 2004, Gamo et al., 2010, Adeyemi et al., 2018, Spalenka et al., 2018). Adapting such high throughput screens to more specific assays offers a route to identifying inhibitors that target particular processes in the parasite. For example, researchers have exploited the observation that the *P. falciparum* ETC becomes dispensable when a cytosolic, CoQ-independent form of DHODH from yeast (yDHODH) is introduced into the parasite (Painter et al., 2007), to develop a more target-based screening approach (Dong et al., 2011). This study identified compounds that have reduced potency against yDHODH-expressing parasites compared to WT *P. falciparum*, and hence are on-target inhibitors of the ETC of these parasites (Dong et al., 2011). Parasite ETC inhibitors have been identified through screening of a compound library using a fluorescence-based Oxygen Biosensor System to directly measure oxygen consumption in erythrocytes infected with *Plasmodium yoelii* (Gomez-Lorenzo et al., 2018). Although this approach is a powerful means of identifying candidate ETC inhibitors, shortcomings of this assay include that it has limited ability to distinguish between on-target ETC inhibitors and off-target compounds that cause parasite death (and therefore lead indirectly to decreased oxygen consumption) (Gomez-Lorenzo et al., 2018), and secondary assays are required to locate the target of identified inhibitors from these screens. An assay in which oxygen consumption and parasite viability could simultaneously be assessed would enable on- and off-target compounds to be differentiated more rigorously, and screening assays that pinpoint the molecular target/s of candidate ETC inhibitors would provide a valuable means of identifying novel targets in the ETC.

Here, we screened the Medicines for Malaria Venture (MMV) ‘Pathogen Box’ small molecule library to identify inhibitors of the *T. gondii* parasite ETC using a Seahorse XFe96 flux analyzer. The Seahorse XFe96 flux analyzer simultaneously measures the oxygen consumption rate (OCR) and extracellular acidification rate (ECAR) of parasites to assess ETC function and general metabolism, respectively, thereby allowing us to distinguish between on- and off-target inhibitors. We identified seven compounds that inhibited *T. gondii* OCR, six of which were on-target ETC inhibitors, and a seventh that simultaneously inhibited ECAR, causing rapid parasite death in an off-target manner. Among these compounds were two chemically novel ETC inhibitors, one of which (MMV688853) was previously characterized as an inhibitor of the parasite calcium dependent protein kinase-1 (CDPK1) protein, and which our data therefore indicate has dual targets. We provide evidence that most of the identified inhibitors are also on-target inhibitors of the *P. falciparum* ETC, illustrating that these compounds have broad utility in targeting this important phylum of parasites. We adapted the Seahorse XFe96 flux analyzer assays to identify the targets of these inhibitors, and determined that most target ETC Complex III in these parasites. We also demonstrate that atovaquone-resistant mutants in both *T. gondii* and *P. falciparum* show limited cross-resistance to some of the identified Complex III inhibitors. Taken together, our work establishes a scalable pipeline to both identify and characterize the targets of inhibitors of the ETC in apicomplexan parasites.

## Results

### Screening the MMV ‘Pathogen Box’ identifies 7 inhibitors of oxygen consumption in *T. gondii*

Apicomplexan parasites require oxygen for one key purpose – to act as the terminal electron acceptor in the mitochondrial ETC. In previous studies, we utilized a Seahorse XFe96 extracellular flux analyzer assay to measure the mitochondrial oxygen consumption rate (OCR) in extracellular tachyzoites (Seidi et al., 2018, Hayward et al., 2021, Hayward et al., 2022). These assays enable the injection of compounds into wells of a 96-well plate prior to measuring parasite OCR, and we demonstrated that injection of the Complex III inhibitor atovaquone rapidly inhibits OCR (Seidi et al., 2018, Hayward et al., 2021). We reasoned that this approach could be used to screen large compound libraries to identify new inhibitors of the parasite ETC. To investigate this, we screened the MMV ‘Pathogen Box’ compound library (a library of ‘diverse, drug-like molecules active against neglected diseases’) for inhibitors of parasite mitochondrial OCR. Of the 400 compounds tested, seven were found to inhibit OCR by more than 30% at 1 μM (Fig. 1).

**Figure 1.**
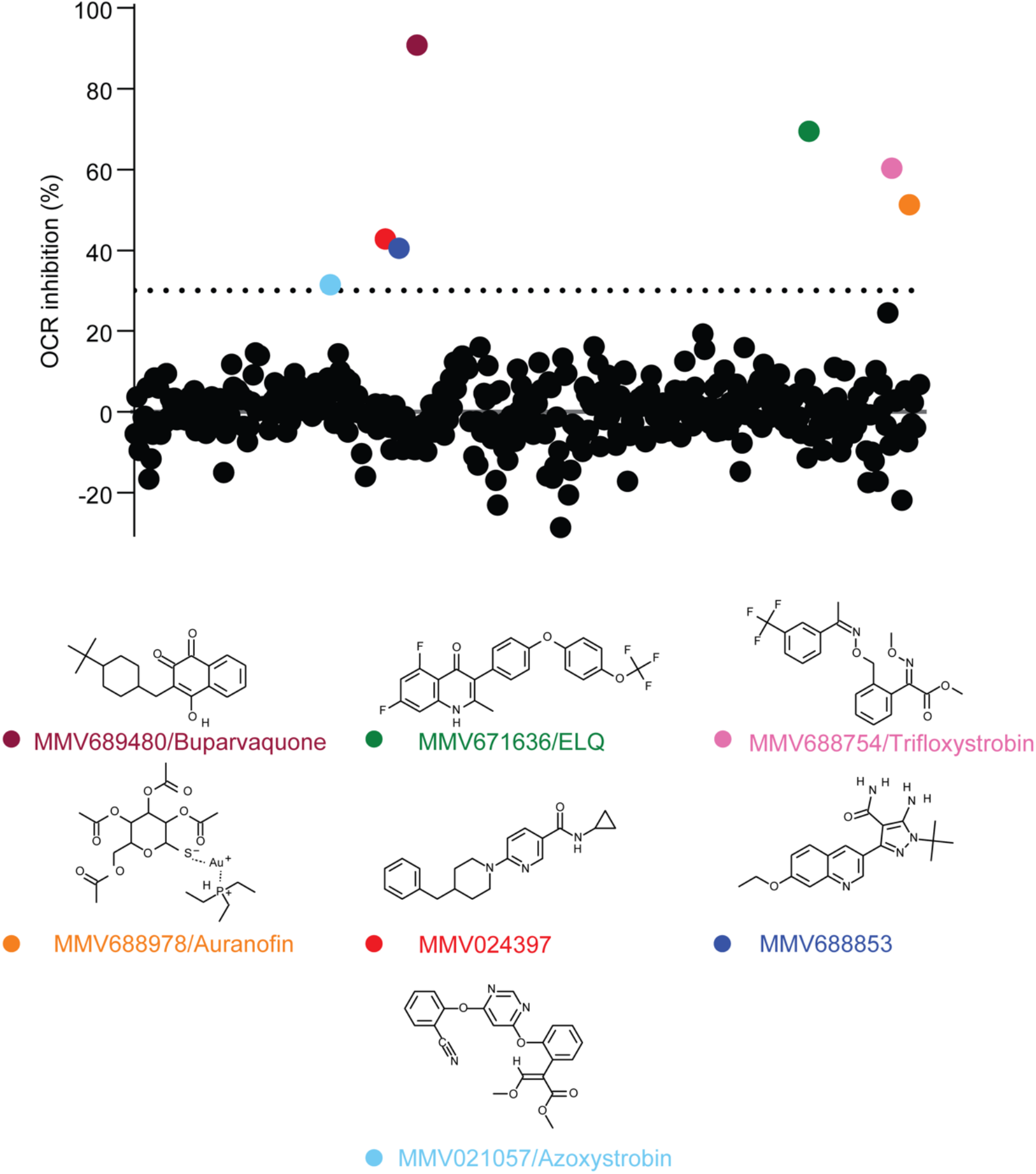
Screening the MMV ‘Pathogen Box’ for inhibitors of O*_2_* consumption in *T. gondii*. The oxygen consumption rate (OCR) of extracellular *T. gondii* parasites was measured in a 96-well plate using a Seahorse XFe96 extracellular flux analyzer. Compounds from the MMV ‘Pathogen Box’ were added to wells at a final concentration of 1 μM, and the change in OCR was monitored in real time after each addition. Percent inhibition of OCR by each of the 400 compounds was calculated relative to complete inhibition observed after addition of 1 µM of the known OCR inhibitor atovaquone, with each compound represented by a dot. A >30% inhibition cut off was applied (dotted line), with seven compounds inhibiting OCR by >30% at 1 μM (coloring of dots corresponds to coloring of labels of the chemical structures shown below). Data are from a single experiment. These hits included MMV689480/buparvaquone (burgundy), the endochin-like quinolone (ELQ) MMV671636 (green), MMV688754/trifloxystrobin (pink), MMV688978/auranofin (orange), MMV024397 (red), the aminopyrazole carboxamide MMV688853 (dark blue), and MMV021057/azoxystrobin (light blue).

Chemically diverse compound scaffolds were represented among the identified hits (Fig. 1), including the known apicomplexan parasite ETC inhibitors MMV689480 (buparvaquone) and the endochin-like quinolone (ELQ) family compound MMV671636. The anti-fungal agents MMV688754 and MMV021057 (trifloxystrobin and azoxystrobin, respectively) were also identified; these compounds bind to the Q_o_ site of Complex III in fungi (Bartlett et al., 2002) and have been shown previously to inhibit *P. falciparum* proliferation (Witschel et al., 2012), likely via binding to the Q_o_ site of Complex III (Vallieres et al., 2013). Other compounds identified in our screen have not yet been shown to be ETC inhibitors and included MMV688853, an aminopyrazole carboxamide compound previously identified as an inhibitor of *T. gondii* calcium-dependent protein kinase 1 (*Tg*CDPK1) (Zhang et al., 2014, Huang et al., 2015), MMV024397 which has been shown to inhibit proliferation of *P. falciparum* (Tougan et al., 2019), and MMV688978 (auranofin). Auranofin is a gold-containing compound used clinically for the treatment of rheumatoid arthritis (Kean et al., 1997), which also inhibits the proliferation of many parasites including *T. gondii* (Ma et al., 2021) and *P. falciparum* (Sannella et al., 2008).

### Identified compounds inhibit proliferation and oxygen consumption in both *T. gondii* and *P. falciparum*

We next tested whether the identified compounds could inhibit proliferation of *T. gondii* parasites. We measured the proliferation of RH strain *T. gondii* tachyzoites expressing a tandem dimeric Tomato (tdTomato) red fluorescent protein using a previously described fluorescence-based 96-well plate proliferation assay (Rajendran et al., 2017). All seven compounds inhibited *T. gondii* proliferation with sub- to high-nanomolar IC_50_ values, with buparvaquone (IC_50_ ± SEM = 0.7 ± 0.1 nM, n = 3) and the ELQ MMV671636 (IC_50_ ± SEM = 3.0 ± 0.2 nM, n = 3) the most potent, and azoxystrobin (IC_50_ ± SEM = 310 ± 32 nM, n = 3) the least (Table 1a; Fig. S1). Given that ELQ compounds are well-characterized ETC inhibitors (Doggett et al., 2012, Stickles et al., 2015), we did not include MMV671636 in further experiments.

**Table 1.** Effects of the identified MMV ‘Pathogen Box’ compounds on *T. gondii* and *P. falciparum* proliferation. **(a)** Determination of the inhibitory properties of the identified compounds on the proliferation of wild type (WT) RH strain, WT ME49 strain, or atovaquone-resistant (ATV^R^) ME49 strain *T. gondii* parasites. **(b)** Determination of the inhibitory properties of the identified compounds on the proliferation of WT 3D7 strain, yeast dihydroorotate dehydrogenase (yDHODH)-expressing 3D7 strain, or ATV^R^ 3D7 strain *P. falciparum* parasites. As the yDHODH and ATV^R^ strains were generated in different laboratories, proliferation of the WT 3D7 background strain of each was determined for comparisons. Data are reported as average IC_50_ (nM) ± SEM from three or more independent experiments. The fold change (FC) was calculated by dividing the IC_50_ against ATV^R^ ME49 parasites by the IC_50_ against WT ME49 *T. gondii* parasites, or the IC_50_ against ATV^R^ 3D7 parasites by the IC_50_ against WT *P. falciparum* parasites, with FC values >1 indicating increased resistance and FC values <1 indicating increased sensitivity of the ATV^R^ strains to the tested compounds. Paired t-tests were performed to compare the IC_50_ of WT and ATV^R^ parasites, and *p*-values are depicted as ns = not significant (*p* > 0.05), * *p* < 0.05, ** *p* < 0.01, *** *p* < 0.001, **** *p* < 0.0001. ND = not determined. NA = not applicable.

| Compound | (a) *T. gondii* |  |  |  | (b) *P. falciparum* |  |  |  |  |
| --- | --- | --- | --- | --- | --- | --- | --- | --- | --- |
|  | RH WT | ME49 WT | ME49 ATV<sup>R</sup> |  | WT | yDHODH | WT | ATV<sup>R</sup> |  |
|  | IC<sub>50</sub> (nM) | IC<sub>50</sub> (nM) | IC<sub>50</sub> (nM) | FC | IC<sub>50</sub> (nM) | IC<sub>50</sub> (nM) | IC<sub>50</sub> (nM) | IC<sub>50</sub> (nM) | FC |
| Atovaquone | 10.4 ± 0.5 | 14 ± 4 | 284 ± 34 | 20 * | 0.13 ± 0.02 | > 10 | 0.31 ± 0.04 | 7.6 ± 0.9 | 24 ** |
| Trifloxystrobin | 28 ± 2 | 67 ± 18 | 24 ± 5 | 0.4 ns | 44 ± 16 | > 250 | 33 ± 7 | 131 ± 12 | 4 * |
| Azoxystrobin | 310 ± 32 | 579 ± 48 | 232 ± 36 | 0.4 * | 23 ± 9 | > 125 | 12 ± 1 | 31 ± 7 | 2.6 ns |
| MMV024397 | 238 ± 30 | 153 ± 18 | 441 ± 92 | 2.9 ns | 308 ± 18 | 3740 ± 1280 | 400 ± 48 | 602 ± 93 | 1.5 * |
| MMV688853 | 69 ± 12 | 178 ± 14 | 133 ± 4 | 0.7 ns | > 6250 | > 6250 | >40 000 | >40 000 | NA |
| Buparvaquone | 0.7 ± 0.1 | 0.7 ± 0.2 | 163 ± 14 | 233 ** | 1.2 ± 0.3 | > 12.5 | 10.9 ± 1.2 | 1160 ± 215 | 106 * |
| Auranofin | 102 ± 27 | 92 ± 13 | 191 ± 44 | 2 ns | 2040 ± 410 | 1810 ± 490 | 2831 ± 503 | 2783 ± 362 | 1 ns |
| MMV671636 | 3.0 ± 0.2 | ND | ND | ND | ND | ND | ND | ND | ND |
| Chloroquine | ND | ND | ND | ND | 6.6 ± 1.3 | 7.70 ± 0.07 | 19 ± 2 | 19 ± 3 | 0 ns |

The ETC is a validated drug target in *P. falciparum* parasites (Barton et al., 2010), and we reasoned that the identified inhibitors of OCR in *T. gondii* may also act against the ETC of *P. falciparum*. We first tested whether the identified compounds could inhibit proliferation of the disease-causing asexual blood stage of 3D7 strain *P. falciparum*. Five of the six compounds inhibited 3D7 *P. falciparum* proliferation, most with sub- to high-nanomolar IC_50_ values (Table 1b; Fig. 2). While MMV688853 was an effective inhibitor of *T. gondii* proliferation, we found that it had little effect on the proliferation of *P. falciparum* at the concentration range we tested (up to 6.25 μM) (Fig. 2h). As an initial measure for whether they act specifically on the ETC of *P. falciparum* or whether they have broader cellular targets, we tested the ability of the identified compounds to inhibit the proliferation of yDHODH-expressing 3D7 strain *P. falciparum* parasites, which are no longer dependent on the ETC for proliferation (Painter et al., 2007). We observed that yDHODH-expressing parasites grew better than WT in the presence of four of the compounds (buparvaquone, trifloxystrobin, azoxystrobin and MMV024397) and the known ETC inhibitor atovaquone (Table 1b; Fig. 2), consistent with these compounds acting primarily on the ETC in *P. falciparum*. By contrast, yDHODH and WT parasites were equally inhibited in the presence of auranofin and the control compound chloroquine, which does not target the ETC (Table 1b; Fig. 2). This observation suggests that auranofin perturbs parasite proliferation independently of ETC inhibition. Together, these results indicate that most of the identified compounds are selective inhibitors of the ETC in *P. falciparum*.

**Figure 2.**
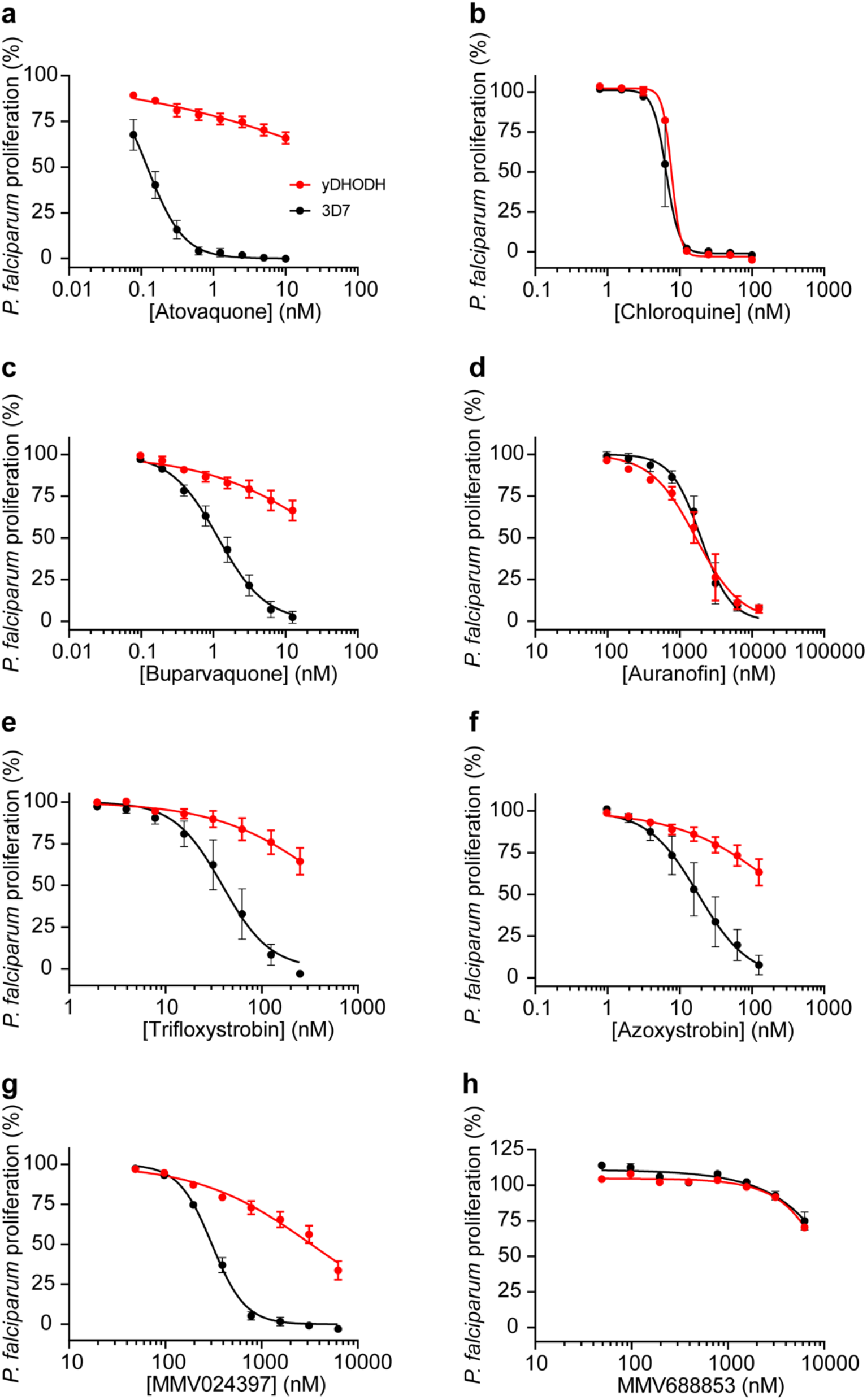
Identification of selective inhibitors of the ETC in *P. falciparum*. Dose-response curves depicting the proliferation of WT (black) or yeast dihydroorotate dehydrogenase (yDHODH)-expressing (red) *P. falciparum* parasites in the presence of increasing concentrations of **(a)** the known ETC inhibitor atovaquone, **(b)** chloroquine, a compound that does not inhibit the ETC, **(c)** buparvaquone, **(d)** auranofin, **(e)** trifloxystrobin, **(f)** azoxystrobin, **(g)** MMV024397, or **(h)** MMV688853 after 96 h of culture. Values are expressed as a percentage of the average proliferation of the drug-free control, and represent the mean ± SEM of three independent experiments performed in triplicate; error bars that are not visible are smaller than the symbol.

To explore their potency at inhibiting OCR in *T. gondii*, we investigated the effects of a range of concentrations of each compound on parasite OCR using the Seahorse XFe96 flux analyzer. All compounds inhibited the OCR of *T. gondii* tachyzoites in a dose-dependent manner (Table 2; Fig. S2c-i). Most of the tested compounds showed rapid inhibition of OCR at the higher concentrations tested (as shown for atovaquone, Fig. S2a). By contrast, inhibition of OCR by auranofin occurred more gradually over time, even at the highest concentration tested (Fig. S2b), suggesting that the effects of auranofin on OCR may occur in a different manner to the other identified compounds.

**Table 2.** Inhibitory activities of MMV ‘Pathogen Box’ compounds against OCR in *T. gondii* and *P. falciparum*. WT *T. gondii* (RH strain) and WT *P. falciparum* (3D7 strain) oxygen consumption rates were assessed using a Seahorse XFe96 flux analyzer. Data are reported as average IC_50_ (μM) ± SEM from three or more independent experiments. ND = not determined.

| Compound | *T. gondii*<br>IC<sub>50</sub> (μM) | *P. falciparum*<br>IC<sub>50</sub> (μM) |
| --- | --- | --- |
| Atovaquone | 0.18 ± 0.05 | 0.022 ± 0.008 |
| Trifloxystrobin | 0.50 ± 0.02 | 0.042 ± 0.017 |
| Azoxystrobin | 7.05 ± 3.08 | 0.015 ± 0.002 |
| MMV024397 | 2.81 ± 0.66 | 0.413 ± 0.051 |
| MMV688853 | 2.76 ± 0.48 | >10 |
| Buparvaquone | 1.18 ± 0.69 | ND |
| Auranofin | 2.48 ± 0.46 | >100 |

In addition to measuring OCR, the Seahorse XFe96 extracellular flux analyzer simultaneously measures the extracellular acidification rate (ECAR), which provides a general measure of parasite metabolic activity (Seidi et al., 2018, Hayward et al., 2021). We observed that most of the test compounds inhibited OCR without inhibiting ECAR (Fig. 3a), suggesting that they selectively target the ETC of the parasite. By contrast, treatment with auranofin resulted in a concomitant decrease in both OCR and ECAR (Fig. 3a). This provides additional evidence that auranofin acts in a different manner to the other identified compounds. To explore this further, we assessed the viability of parasites upon auranofin treatment. We treated *T. gondii* parasites with 1, 20 or 100 µM auranofin, or 10 µM atovaquone as a control, stained parasites with propidium iodide (PI), and quantified parasite viability by flow cytometry. We observed that treatment with auranofin led to a rapid, dose-dependent decrease in parasite viability over the 140-minute time course of the assay (Fig. 3b). By contrast, treatment with the selective ETC inhibitor atovaquone caused minimal loss of parasite viability within this timeframe (Fig. 3b), suggesting the decreased viability observed upon auranofin treatment is not due to ETC inhibition. These data suggest that auranofin is not a selective inhibitor of the ETC but instead perturbs broader parasite metabolic functions, resulting in a decrease in parasite viability and a secondary impairment of ETC activity.

**Figure 3.**
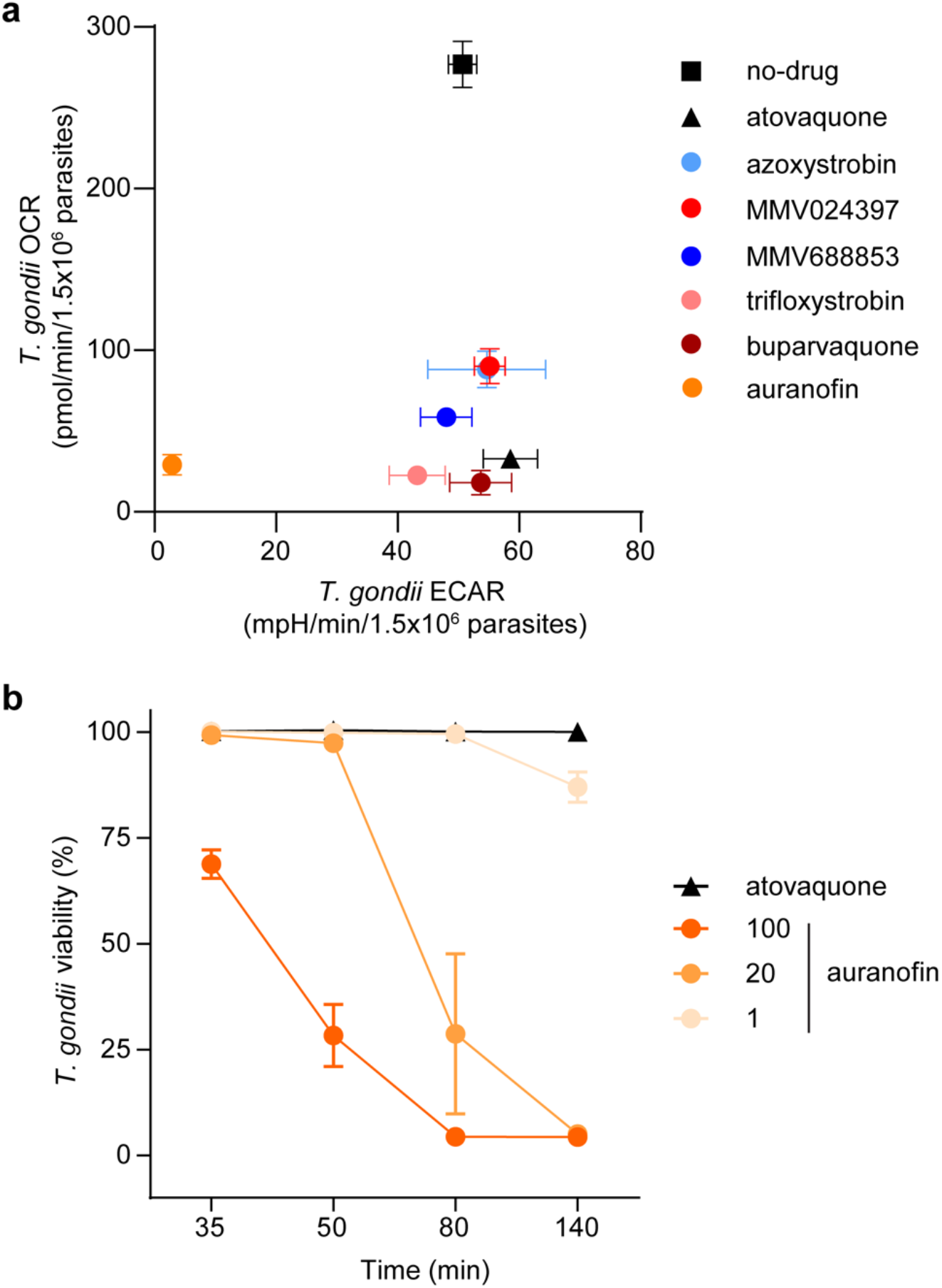
Identification of selective and off-target inhibitors of the ETC in *T. gondii* parasites. **(a)** Oxygen consumption rate (OCR) versus extracellular acidification rate (ECAR) of *T. gondii* parasites treated with either no-drug (black square), atovaquone (black triangle; 10 μM), azoxystrobin (light blue; 80 μM), MMV024397 (red; 20 μM), MMV688853 (dark blue; 20 μM), trifloxystrobin (pink; 10 μM), buparvaquone (burgundy; 20 μM) or auranofin (orange; 80 μM) assessed using a Seahorse XFe96 flux analyzer. Data represent the mean OCR and ECAR ± SEM of three independent experiments, and are derived from the top concentration of inhibitor tested in Fig. S2. **(b)** Viability of extracellular *T. gondii* parasites treated with atovaquone (black triangles, 10 μM) or auranofin (orange circles, 1-100 μM) for 35 – 140 minutes. Viability was assessed by flow cytometry of propidium iodide-stained parasites and normalized to a DMSO-treated vehicle control. Data represent the mean ± SEM of three independent experiments; error bars that are not visible are smaller than the symbol.

We conclude that most of the compounds identified in our initial screen inhibit the proliferation of *T. gondii* and *P. falciparum* parasites, and act selectively on the ETC of these parasites. A strength of the Seahorse XFe96 flux analyzer-based screening approach is its ability to simultaneously measure OCR and ECAR, and thereby enable the differentiation of compounds that directly inhibit the ETC from those – such as auranofin – that have a broader effect on parasite metabolism or viability.

### MMV688853 inhibits the ETC in a *Tg*CDPK1-independent manner

One of the hit compounds identified in our ETC inhibitor screen was the aminopyrazole carboxamide scaffold compound MMV688853, which has been reported previously to be an inhibitor of *T. gondii* calcium-dependent protein kinase 1 (*Tg*CDPK1) (Zhang et al., 2014, Huang et al., 2015). *Tg*CDPK1 is a cytosolic protein that has been shown to be critical for parasite invasion of host cells (Lourido et al., 2010). We hypothesized that either *Tg*CDPK1 has an additional role in the ETC or that MMV688853 has a second target in these parasites. *Tg*CDPK1 has a glycine residue at the mouth of the pocket where MMV688853 and other *Tg*CDPK1 inhibitors bind (Fig. 4a). Mutation of this so-called ‘gatekeeper’ residue to a bulky amino acid like methionine renders *Tg*CDPK1 resistant to inhibition by aminopyrazole carboxamide scaffold compounds like MMV688853 (Huang et al., 2015), as well as to pyrazolopyrimidine scaffold compounds such as 3-MB-PP1 (Lourido et al., 2010) (Fig. 4a). To test our hypotheses, we generated a tdTomato^+^ *T. gondii* strain wherein the gatekeeper glycine residue at position 128 was mutated to methionine (*Tg*CDPK1^G128M^; Fig. 4a).

**Figure 4.**
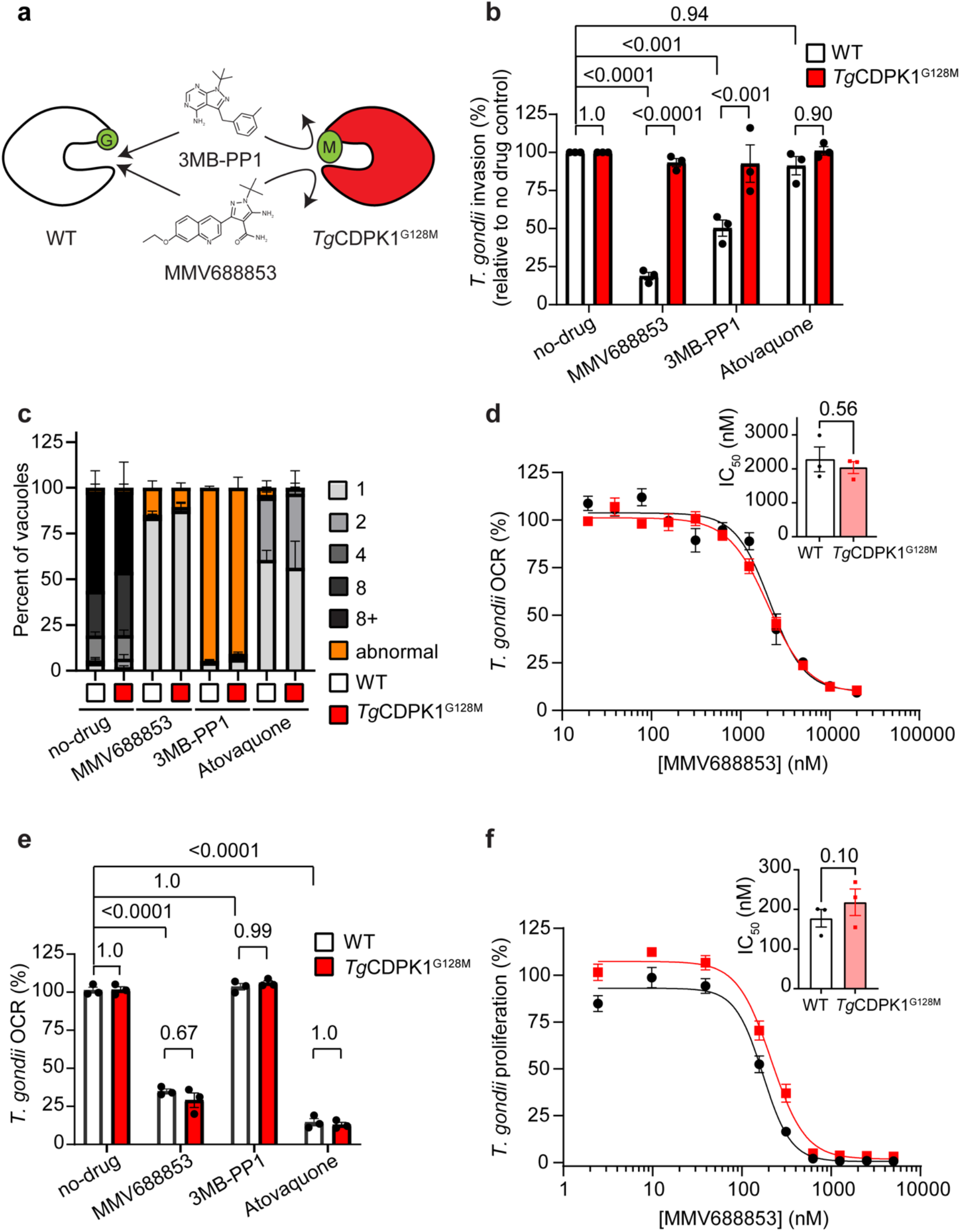
MMV688853 dually targets *Tg*CDPK1 and the ETC in *T. gondii* parasites. **(a)** Schematic depicting the small glycine gatekeeper residue of WT *Tg*CDPK1 (white) which enables inhibition by 3MB-PP1 and MMV688853. Mutation of this residue to a larger methionine (*Tg*CDPK1^G128M^, red) blocks inhibitor access to the binding site and thereby confers resistance to these compounds. **(b)** Percent invasion of parasites expressing WT *Tg*CDPK1 (white) or *Tg*CDPK1^G128M^ (red) into host cells in the absence of drug (DMSO vehicle control), or the presence of MMV688853 (5 μM), 3MB-PP1 (5 μM) or atovaquone (1 μM), normalized relative to the no-drug control. At least 100 parasites were counted per experiment, with data representing the mean ± SEM of three independent experiments (each experiment shown as a dot). ANOVA followed by Tukey’s multiple comparisons test was performed with relevant *p*-values shown. **(c)** Intracellular proliferation assays depicting the percent of vacuoles containing 1-8+ (gray tones) or abnormal (orange) parasites when parasites expressing WT *Tg*CDPK1 (white) or *Tg*CDPK1^G128M^ (red) were cultured in the absence of drug (DMSO vehicle control), or the presence of MMV688853 (5 μM), 3MB-PP1 (5 μM) or atovaquone (1 μM) for 20 h. Abnormal morphology was defined as vacuoles that contained misshapen parasites. At least 100 vacuoles were counted per condition, with data representing the mean ± SEM of three independent experiments. **(d)** Dose-response curves depicting the oxygen consumption rate (OCR) of parasites expressing WT *Tg*CDPK1 (black) or *Tg*CDPK1^G128M^ (red) incubated with increasing concentrations of MMV688853 as a percentage of a no-drug (DMSO vehicle) control. Data represent the mean ± SEM of three independent experiments. Inset bar graph depicts the IC_50_ ± SEM (nM) of three independent experiments (each experiment shown as a dot). The *p*-value from a paired t-test is shown. **(e)** OCR of parasites expressing WT *Tg*CDPK1 (white) or *Tg*CDPK1^G128M^ (red) incubated in the absence of drug (DMSO vehicle control), or in the presence of MMV688853 (5 μM), 3MB-PP1 (5 μM) or atovaquone (1 μM), expressed as a percentage of the OCR prior to addition of compounds. Data represent the mean ± SEM of three independent experiments. ANOVA followed by Tukey’s multiple comparisons test was performed with relevant *p*-values shown. **(f)** Dose-response curves depicting the percentage proliferation of parasites expressing WT *Tg*CDPK1 (black) or *Tg*CDPK1^G128M^ (red) in the presence of increasing concentrations of MMV688853 over 6 days. Values are expressed as a percent of the average fluorescence from the no-drug control at mid-log phase growth in the fluorescence proliferation assay, and represent the mean ± SEM of three independent experiments; error bars that are not visible are smaller than the symbol. Inset bar graph depicts the IC_50_ ± SEM (nM) of three independent experiments (each experiment shown as a dot). The *p*-value from a paired t-test is shown.

*Tg*CDPK1 is an important regulator of parasite invasion (Lourido et al., 2010), a critical step in the lytic cycle of the parasite. Previous studies have shown that *Tg*CDPK1 inhibitors impair host cell invasion by WT but not *Tg*CDPK1^G128M^ parasites (Lourido et al., 2010). To validate this, we tested the ability of MMV688853 to inhibit the invasion of WT and *Tg*CDPK1^G128M^ parasites. While invasion of WT parasites was significantly inhibited by both MMV688853 and the control *Tg*CDPK1 inhibitor 3-MB-PP1, *Tg*CDPK1^G128M^ parasites were able to invade in the presence of either compound (Fig. 4b). By comparison, the ETC inhibitor atovaquone did not inhibit the invasion of either parasite strain (Fig. 4b). These results indicate that MMV688853 inhibits *T. gondii* invasion in a *Tg*CDPK1-dependent manner.

We next tested the ability of MMV688853 to inhibit intracellular proliferation of WT and *Tg*CDPK1^G128M^ parasites. We allowed parasites to invade host cells in the absence of inhibitors, then grew parasites for ∼20 h in the presence of MMV688853 or various control inhibitors and quantified the number of parasites per vacuole. MMV688853 inhibited intracellular proliferation of both WT and *Tg*CDPK1^G128M^ parasites, with most vacuoles having only a single parasite (Fig. 4c). Treatment with atovaquone resulted in similar impairment of intracellular proliferation (Fig. 4c), with the majority of vacuoles containing 1-2 parasites. These data indicate that MMV688853 can inhibit intracellular proliferation independently of *Tg*CDPK1. Unexpectedly, the majority of both WT and *Tg*CDPK1^G128M^ parasites grown in the presence of 3-MB-PP1 exhibited abnormal morphology (defined as vacuoles that contained misshapen parasites, possibly resulting from defects in cell division; Fig. 4c), suggesting an additional off-target effect of 3-MB-PP1.

As a test for whether the inhibition of oxygen consumption by MMV688853 occurs through inhibition of *Tg*CDPK1, we assessed the OCR of intact WT and *Tg*CDPK1^G128M^ parasites after addition of increasing concentrations of MMV688853. We observed a similar IC_50_ for OCR inhibition in both WT and *Tg*CDPK1^G128M^ parasites (Fig. 4d). We also examined the ability of the alternative *Tg*CDPK1 inhibitor 3-MB-PP1 to inhibit OCR of WT and *Tg*CDPK1^G128M^ parasites. In contrast to atovaquone and MMV688853, 3-MB-PP1 did not inhibit OCR in either WT or *Tg*CDPK1^G128M^ parasites (Fig. 4e). Together, these data indicate that MMV688853 acts on the ETC independently of *Tg*CDPK1, and that *Tg*CDPK1 does not have a role in the ETC.

Finally, we measured the effects of MMV688853 on the overall proliferation of WT and *Tg*CDPK1^G128M^ *T. gondii* parasites through the lytic cycle. We measured parasite proliferation in the presence of increasing concentrations of MMV688853 over six days using a fluorescence proliferation assay. We observed a similar IC_50_ for both WT and *Tg*CDPK1^G128M^ parasites (Fig. 4f). Taken together, our data indicate that while MMV688853 inhibits parasite invasion by targeting *Tg*CDPK1 (Fig. 4b), MMV688853 also has a second target in the ETC of the parasite (Fig. 4d-e), and this second target is likely a major contributor to impairment of intracellular proliferation of the parasite by this compound (Fig. 4f).

### Defining the targets of the candidate ETC inhibitors in *T. gondii* and *P. falciparum*

Having characterized the inhibitory properties of the candidate ETC inhibitors, we next sought to identify which component of the ETC these compounds target. To do this, we utilized a Seahorse XFe96 analyzer-based assay that we developed previously to pinpoint where a defect in the *T. gondii* ETC is occurring (Hayward et al., 2021, Hayward et al., 2022) (Fig. 5a). Briefly, *T. gondii* parasites were starved for 1 h to deplete endogenous substrates. The plasma membrane of the parasites was permeabilized using a low concentration of the detergent digitonin, and parasites were incubated with one of two substrates that independently feed electrons to CoQ in the mitochondrion: 1) malate, which donates electrons to the ETC via a reaction catalyzed by the TCA cycle enzyme malate:quinone oxidoreductase; or 2) glycerol 3-phosphate (G3P), which donates electrons to the ETC independently of the TCA cycle via a reaction catalyzed by G3P dehydrogenase. Following substrate addition, the candidate inhibitor was added at a concentration that we previously showed completely inhibited OCR (Fig. S2) and the change in OCR was measured. If OCR elicited by both substrates was inhibited, this provided evidence that the inhibitor was acting downstream of CoQ (*i.e.* on ETC Complexes III or IV) (Hayward et al., 2021) (Fig. 5a). To differentiate between Complex III and Complex IV inhibition, samples were next treated with the substrate *N*,*N*,*N*´,*N*´-tetramethyl-*p*-phenylenediamine dihydrochloride (TMPD), which donates electrons directly to CytC and consequently bypasses Complex III (Fig. 5a). If inhibition of OCR was rescued by addition of TMPD, this provided evidence that the inhibitor was acting upstream of CytC (*e.g.* on ETC Complex III). Finally, samples were treated with the Complex IV inhibitor sodium azide (NaN_3_) to validate that the observed TMPD-dependent OCR was a result of Complex IV activity.

**Figure 5.**
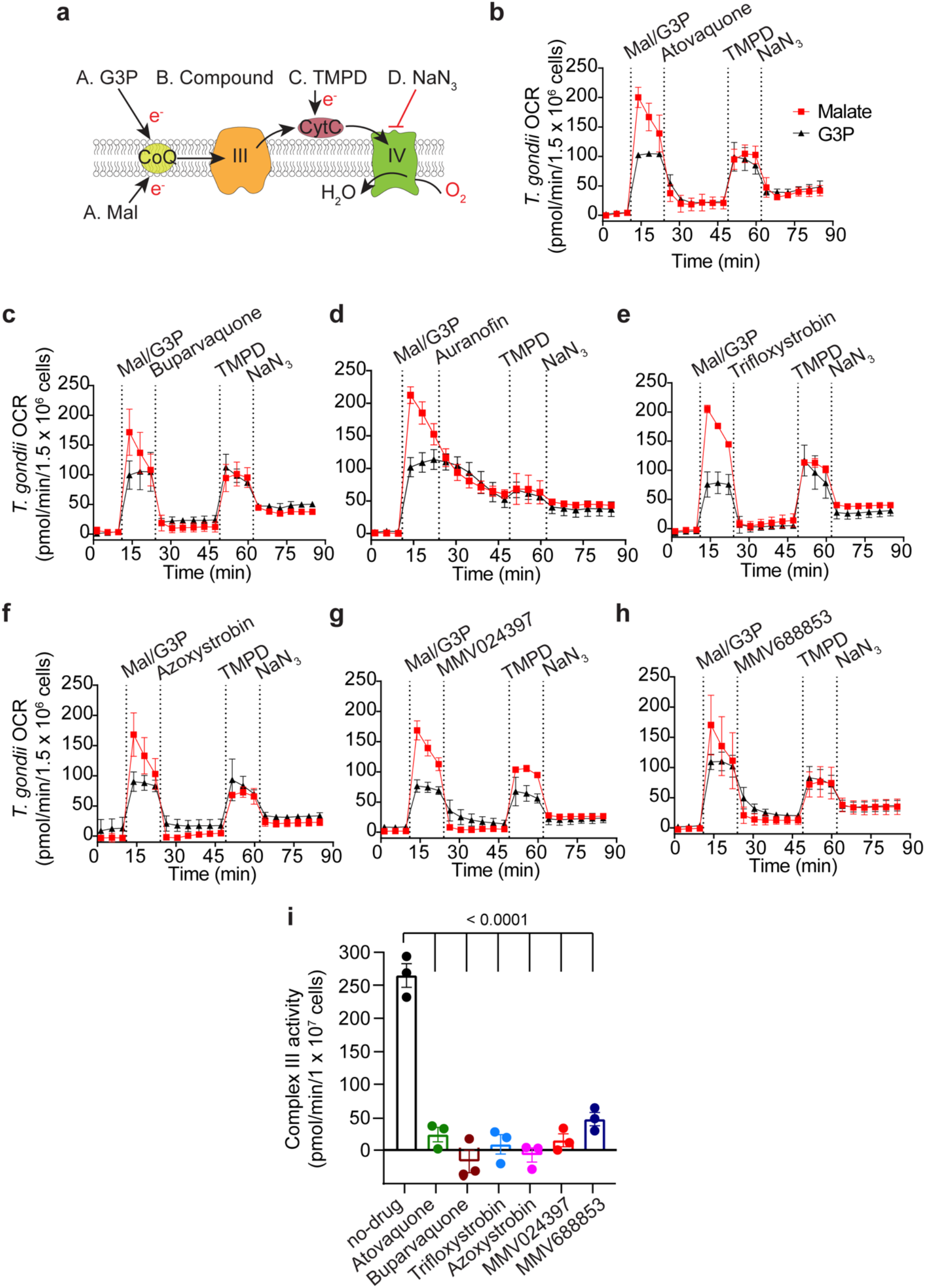
An assay to characterize the targets of the candidate ETC inhibitors identifies chemically diverse Complex III inhibitors. **(a)** Schematic of the assay measuring the oxygen consumption rate (OCR) of plasma membrane-permeabilized *T. gondii* parasites. Parasites were starved for 1 hour to deplete endogenous substrates then permeabilized with digitonin before the addition of the following substrates and inhibitors: Port A, the substrates malate (Mal) or glycerol 3-phosphate (G3P); Port B, the test compound; Port C, TMPD; Port D, sodium azide (NaN_3_). CoQ, coenzyme Q; III, Complex III; CytC, cytochrome *c*; IV, Complex IV; e^-^, electrons. **(b-h)** Traces depicting parasite OCR over time when supplying Mal (red squares) or G3P (black triangles) as a substrate. The candidate ETC inhibitors were **(b)** atovaquone (1.25 μM), **(c)** buparvaquone (5 μM), **(d)** auranofin (10 μM), **(e)** trifloxystrobin (2.5 μM), **(f)** azoxystrobin (80 μM), **(g)** MMV024397 (20 μM), **(h)** MMV688853 (20 μM). Values represent the mean ± SD of three technical replicates and are representative of three independent experiments; error bars that are not visible are smaller than the symbol. **(i)** *T. gondii* Complex III enzymatic activity was assessed in the presence of DMSO (no-drug), atovaquone (1.25 μM), buparvaquone (5 μM), trifloxystrobin (2.5 μM), azoxystrobin (80 μM), MMV024397 (20 μM) or MMV688853 (20 μM). Data represent the mean ± SEM of three independent experiments each conducted in duplicate, with the mean of each experiment represented by a dot. ANOVA followed by Dunnett’s multiple comparisons test were performed and *p*-values are shown.

We observed that most compounds inhibited OCR regardless of whether the parasites were utilizing malate or G3P as ETC substrates (Fig. 5b-h), suggesting that inhibition by these compounds was occurring downstream of CoQ. While most compounds inhibited OCR almost immediately, auranofin inhibition was more gradual (Fig. 5d), consistent with our previous evidence of indirect inhibition of the ETC by this compound (Fig. 3). Furthermore, OCR could be rescued by TMPD for all compounds except auranofin (Fig. 5b-h), which indicates that these compounds inhibit upstream of CytC. Together, these data indicate that the on-target compounds identified in our screen all act via inhibition of ETC Complex III.

To validate these results, we performed a direct, spectrophotometric-based Complex III enzymatic assay on parasite extracts in the absence or presence of inhibitors. We observed that Complex III activity was significantly lower in the presence of all tested on-target inhibitors than in the no-drug control (Fig. 5i; Fig. S3), suggesting that the identified compounds are indeed Complex III inhibitors. The inhibitory activity of auranofin could not be assessed via this assay since we observed apparent enzyme activity upon auranofin addition even in the absence of parasite extract (Fig. S3a).

To begin to define the targets of the identified compounds in *P. falciparum*, we tested the ability of the compounds to inhibit OCR in permeabilized *P. falciparum* parasites that were supplied malate as a substrate (Fig. 6a). We observed that all compounds except auranofin (Fig. 6c) and MMV688853 (Fig. 6g) could inhibit OCR of *P. falciparum* parasites, and that TMPD restored OCR in all cases (Fig. 6b-g). These results are consistent with most of the compounds that inhibited Complex III in *T. gondii* inhibiting the same complex in *P. falciparum*, although our assay cannot rule that they targeting malate oxidation instead. To investigate the potency of each compound in inhibiting OCR of *P. falciparum*, we performed a dose-response experiment (Fig. S4). All compounds except MMV688853 (Fig. S4f) and auranofin (Fig. S4b) inhibited OCR of digitonin-permeabilized *P. falciparum* in a dose-dependent manner, with IC_50_ values in the sub-micromolar range (Table 2). Together, our data provide evidence that most identified compounds are potent inhibitors of the ETC in both *T. gondii* and *P. falciparum*, and that these compounds likely target Complex III. Our data also point to some differences in the activity of these compounds between *T. gondii* and *P. falciparum*, most notably with MMV688853, which inhibits Complex III in *T. gondii*, but is inactive against the ETC in *P. falciparum* at the concentrations tested.

**Figure 6.**
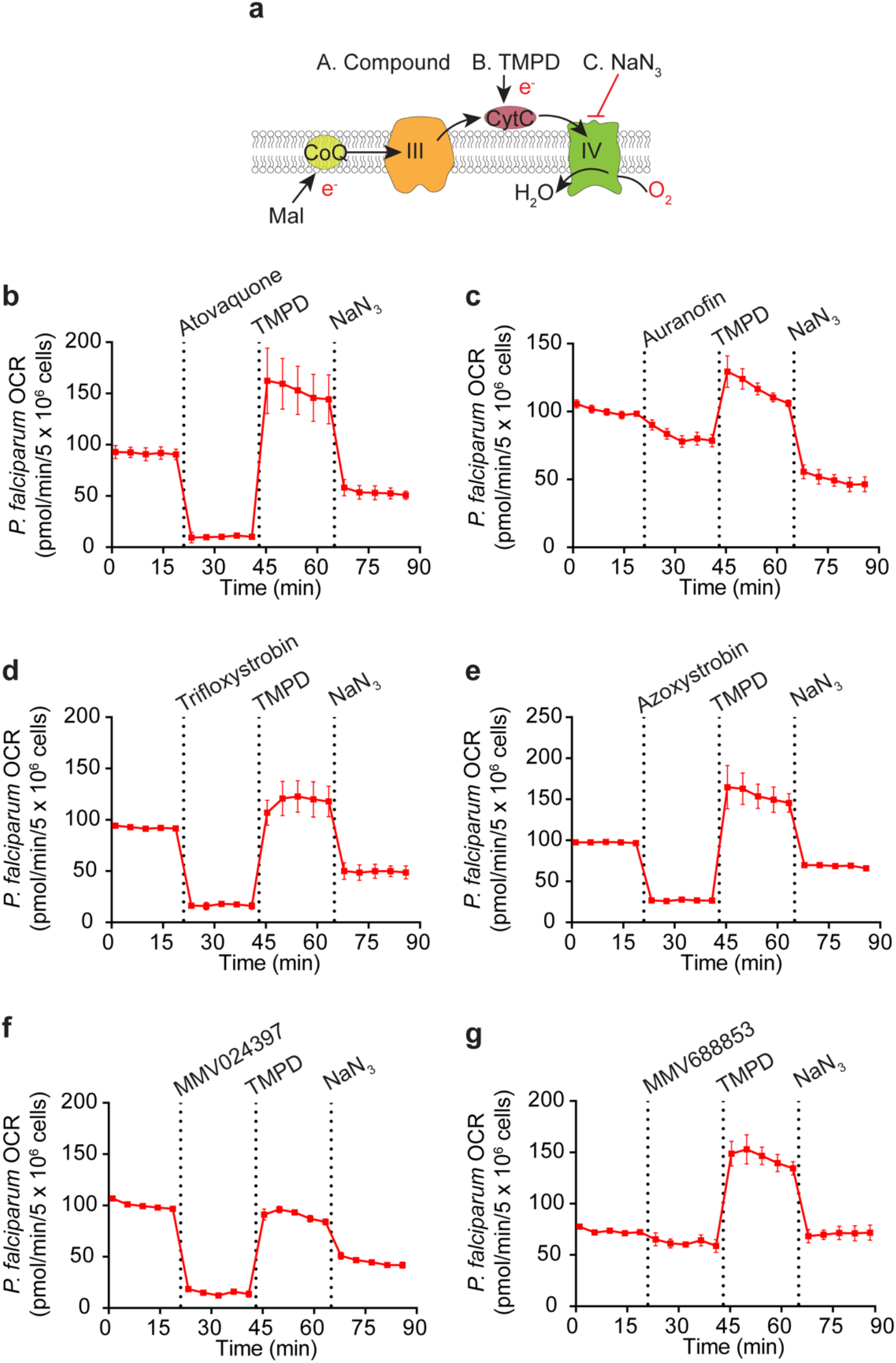
Most of the candidate ETC inhibitors target the ETC upstream of cytochrome *c* in *P. falciparum* parasites. **(a)** Schematic of the assay measuring the oxygen consumption rate (OCR) of permeabilized *P. falciparum* parasites supplied malate (Mal) as a substrate. The following addition of substrates and inhibitors were performed: Port A, the test compound; Port B, TMPD; Port C, sodium azide (NaN_3_). CoQ, coenzyme Q; III, Complex III; CytC, cytochrome *c*; IV, Complex IV; e^-^, electrons. **(b-g)** Traces depicting parasite OCR over time when supplying Mal as a substrate. The candidate ETC inhibitors tested (all at 10 μM) were **(b)** atovaquone, **(c)** auranofin, **(d)** trifloxystrobin, **(e)** azoxystrobin, **(f)** MMV024397, **(g)** MMV688853. Values represent the mean ± SD of three technical replicates and are representative of three independent experiments; error bars that are not visible are smaller than the symbol.

### Atovaquone-resistant *T. gondii* and *P. falciparum* exhibit limited cross-resistance to most of the identified MMV compounds

Atovaquone resistance is known to arise rapidly in apicomplexans, both in the field and the laboratory (Looareesuwan et al., 1996, Cottrell et al., 2014, McFadden et al., 2000). Atovaquone acts by binding the Q_o_ CoQ binding site of Complex III, which is a pocket formed by the cytochrome *b* protein of Complex III. Mutations in Q_o_ site residues of cytochrome *b*, a gene encoded on the mitochondrial genome of apicomplexan parasites, confer varying degrees of atovaquone resistance in both *T. gondii* and *Plasmodium spp.* (McFadden et al., 2000, Srivastava et al., 1999, Syafruddin et al., 1999). We tested whether atovaquone-resistant strains of *T. gondii* and *P. falciparum* parasites exhibited cross-resistance to any of the Complex III inhibitors identified in our screen.

We first tested the effects of the identified inhibitors on a previously described atovaquone-resistant (ATV^R^) ME49 strain of *T. gondii* which has an isoleucine to leucine substitution at position 254 (I254L) of cytochrome *b* (McFadden et al., 2000). We integrated tdTomato into WT (ME49 WT) and ATV^R^ *T. gondii* and performed fluorescence proliferation assays to compare the ability of atovaquone and the test compounds to inhibit proliferation of these two strains. As demonstrated for RH strain parasites (Table 1), all six compounds inhibited WT ME49 strain *T. gondii* proliferation at sub-micromolar concentrations (Table 1a; Fig. 7). As expected, the ATV^R^ strain was resistant to atovaquone, with a ∼20-fold higher IC_50_ than WT parasites (*p* = 0.017; Table 1 a; Fig. 7a). ATV^R^ parasites were cross-resistant to buparvaquone (∼233-fold, *p* = 0.0076; Table 1 a; Fig. 7b). Interestingly, ATV^R^ *T. gondii* parasites were slightly sensitized to the antifungal strobilurin family compounds azoxystrobin (∼2.5-fold, *p* = 0.025; Table 1a; Fig. 7e) and trifloxystrobin (∼2.5-fold, *p* = 0.077; Table 1a; Fig. 7d), and showed minimal cross-resistance against the other tested inhibitors.

**Figure 7.**
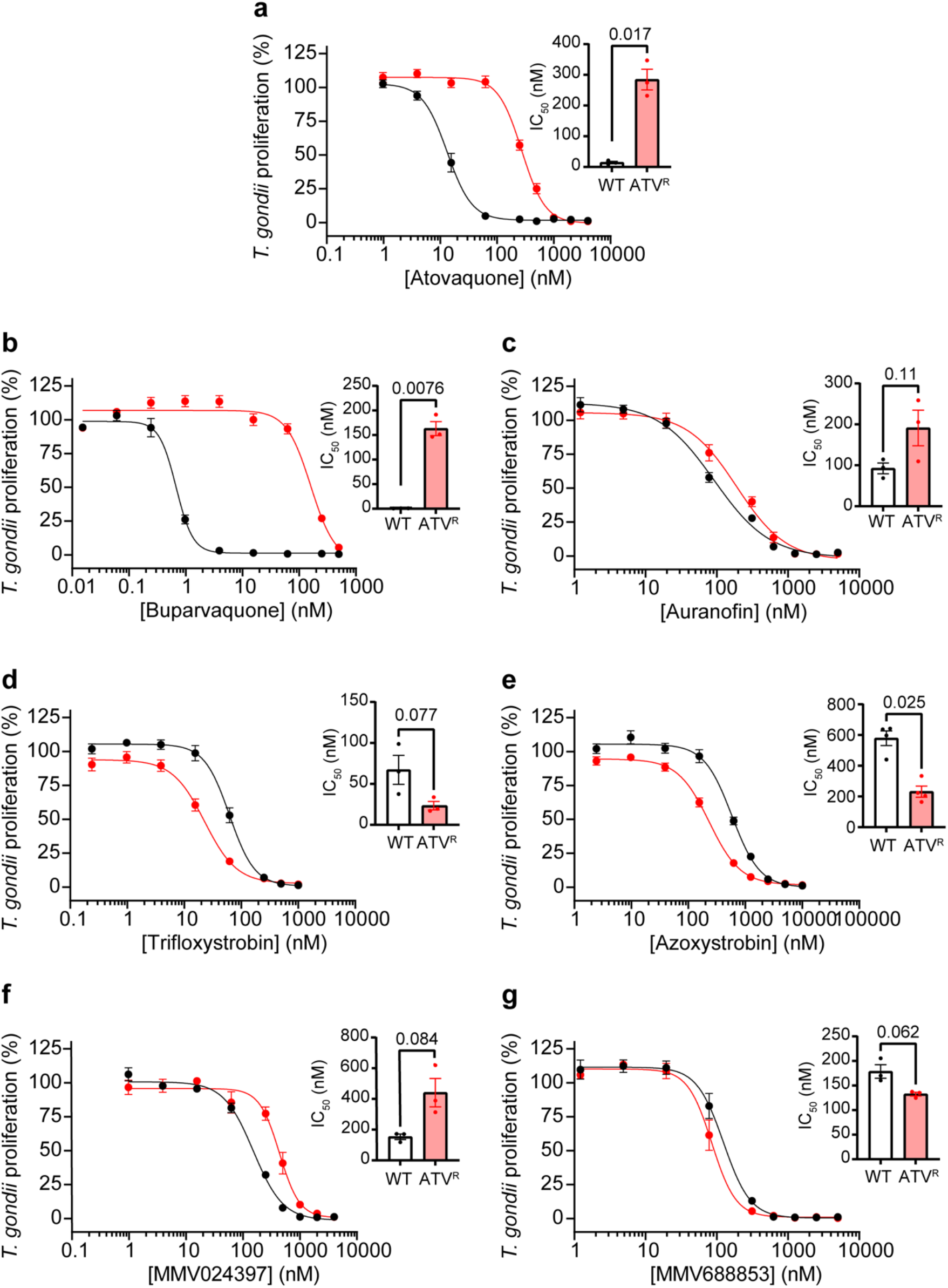
Assessing the activity of ETC inhibitors against atovaquone-resistant *T. gondii* parasites. **(a-g)** Dose-response curves depicting the percent proliferation of WT (black) or atovaquone-resistant (ATV^R^, red) *T. gondii* parasites in the presence of increasing concentrations of **(a)** atovaquone, **(b)** buparvaquone, **(c)** auranofin, **(d)** trifloxystrobin, **(e)** azoxystrobin, **(f)** MMV024397, or **(g)** MMV688853. Values are expressed as a percent of the average fluorescence from a no-drug control at mid-log phase growth in the fluorescence proliferation assay, and represent the mean ± SEM of three (or four for (e)) independent experiments performed in triplicate; error bars that are not visible are smaller than the symbol. Inset bar graphs depict the IC_50_ ± SEM (nM) of three (or four for (e)) independent experiments, with each repeat shown as a dot. Paired t-tests were performed and *p*-values are shown.

We next tested whether an atovaquone resistance-conferring mutation in *P. falciparum* would result in similar changes in sensitivity to the inhibitors identified from our screen. We generated an atovaquone-resistant (ATV^R^) *P. falciparum* parasite strain by drug pressure which had a valine to leucine substitution at position 259 (V259L) in cytochrome *b,* and compared their proliferation in the presence of the candidate ETC inhibitors to WT parasites (Fig. 8). As expected, the ATV^R^ *P. falciparum* strain was resistant to atovaquone, with ∼24-fold higher IC_50_ than WT parasites (*p* = 0.0087; Table 1b; Fig. 8a), but not to chloroquine (Table 1b; Fig. 8b). Like in *T. gondii*, we observed cross-resistance to buparvaquone (∼106-fold, *p* = 0.017; Table 1b; Fig. 8c). We observed little to no cross-resistance of ATV^R^ *P. falciparum* parasites to auranofin (no change; Table 1b; Fig. 8d), trifloxystrobin (∼4-fold, *p* = 0.012; Table 1b; Fig. 8e), azoxystrobin (∼1.5 fold, *p* = 0.055; Table 1b; Fig. 8f), or MMV024397 (∼1.5 fold, *p* = 0.028; Table 1b; Fig. 8g). MMV688853 exhibited minimal inhibition of parasite proliferation in the ATV^R^ strain even at the highest concentration tested (40 µM; Table 1b; Fig. 8h), consistent with the previous assays with WT *P. falciparum* (Fig. 2h). Together, these data indicate that ATV^R^ parasites do not exhibit a great degree of cross-resistance to most of our compounds (with the exception of buparvaquone, which belongs to the same hydroxy-naphthoquinone class as atovaquone).

**Figure 8:**
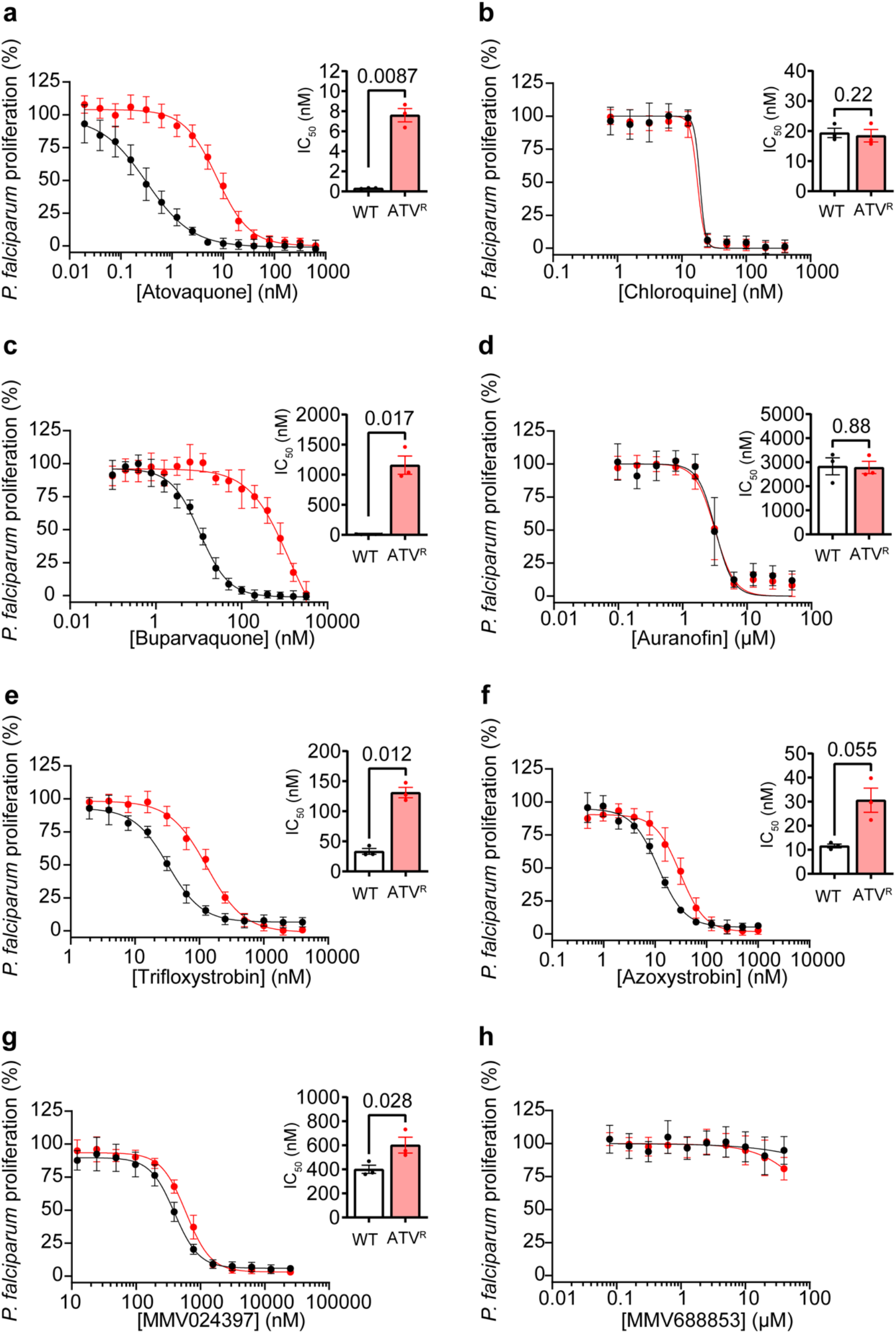
Assessing the activity of ETC inhibitors against atovaquone-resistant *P. falciparum* parasites. **(a-g)** Dose-response curves depicting the percent proliferation of WT (black) or atovaquone-resistant (ATV^R^, red) *P. falciparum* parasites in the presence of increasing concentrations of **(a)** atovaquone, **(b)** chloroquine, **(c)** buparvaquone, **(d)** auranofin, **(e)** trifloxystrobin, **(f)** azoxystrobin, **(g)** MMV024397, or **(h)** MMV688853 after 96 h of culture, as measured using the SYBR Safe-based growth assay. Values are expressed as a percent of the average fluorescence from the no-drug control, and represent the mean ± SEM of three independent experiments performed in triplicate; error bars that are not visible are smaller than the symbol. Inset bar graphs depict the IC_50_ ± SEM (nM) of three independent experiments, with each repeat shown as a dot. Paired t-tests were performed and *p*-values are shown.

## Discussion

In this study, we screened the MMV ‘Pathogen Box’ compound library to identify inhibitors of the *T. gondii* ETC using a Seahorse XFe96 flux analyzer (Fig. 1). One key benefit of using the Seahorse XFe96 flux analyzer as a drug-screening platform is that it simultaneously measures the oxygen consumption rate (OCR) and extracellular acidification rate (ECAR) of parasites to assess ETC function and general metabolism, respectively. This enables on-target ETC inhibitors (*i.e.* those that inhibit OCR but not ECAR) to be differentiated from off-target compounds wherein the defect in OCR is a secondary effect resulting from rapid parasite death or otherwise impaired parasite metabolism (*i.e.* those that inhibit both OCR and ECAR). This is exemplified by the compound auranofin, which inhibited both OCR and ECAR of *T. gondii* and was subsequently shown to induce rapid parasite death (Fig. 3). Furthermore, auranofin inhibited the proliferation of WT and yDHODH-expressing *P. falciparum* to a similar extent (Fig. 2), providing additional evidence that auranofin is unlikely to kill apicomplexan parasites via a direct effect on the ETC. Auranofin has been recently linked to the production of reactive oxygen species (ROS) in *T. gondii* (Ma et al., 2021). Mitochondrial ROS can lead to impairment of ETC function in other organisms (Paradies et al., 2000), which could explain the effects of auranofin on the ETC of *T. gondii*.

Another benefit of the screening approach that we have established is its scalability. By injecting three test compounds sequentially into each well, we were able to screen the entire 400 compound MMV ‘Pathogen Box’ using two 96-well Seahorse XFe96 plates. We note that it is possible to screen much larger compound libraries using this approach.

In addition to compound identification, our approach enables a determination of where in the ETC identified inhibitors target. Using an assay to pinpoint the location of ETC defects in *T. gondii* (Hayward et al., 2021, Hayward et al., 2022), we demonstrated that most compounds identified in our screen (with the exception of auranofin) likely target ETC Complex III (Fig. 5). Specifically, we demonstrated that: 1) the identified compounds inhibited OCR regardless of the electron source (malate or glycerol 3-phosphate) that was donating electrons to CoQ, implying that the inhibition occurred downstream of CoQ; and 2) a substrate that donates electrons directly to CytC (TMPD), and thereby bypasses Complex III, restored OCR, implying that the inhibition occurred upstream of CytC. The druggability of ETC Complex III in apicomplexan parasites has been noted before (Barton et al., 2010). For instance, all seven novel hits identified in a screen for *Plasmodium* ETC inhibitors were found to target Complex III (Gomez-Lorenzo et al., 2018). Our data do not rule out the possibility that, in addition to inhibition of Complex III, the identified compounds also inhibit targets upstream in the ETC (*e.g.* one or more of the dehydrogenases that donate electrons to coenzyme Q). For instance, while the ETC inhibitor 1-hydroxy-2-dodecyl-4(*1H*)quinolone can target Complex III, it can also inhibit DHODH and the single subunit NADH dehydrogenases of apicomplexan parasites (Saleh et al., 2007, Vallieres et al., 2012, Hegewald et al., 2013, Ke et al., 2019), likely by binding the CoQ binding sites of each.

*P. falciparum* rapidly develops resistance to the Complex III inhibitor atovaquone when used in a clinical setting (Looareesuwan et al., 1996, Cottrell et al., 2014), and although atovaquone-resistant clinical isolates of *T. gondii* have not been observed, patients treated with atovaquone frequently experience reactivation of toxoplasmosis (Winterhalter et al., 2010, Chirgwin et al., 2002, Baatz et al., 2006). Atovaquone resistance arises from mutations in the Q_o_ site of the cytochrome *b* protein of Complex III (Vaidya et al., 1993, Srivastava et al., 1999, Syafruddin et al., 1999, McFadden et al., 2000). We tested our identified inhibitors against atovaquone-resistant strains of both *T. gondii* and *P. falciparum* (Fig. 7 and 8). We found that ATV^R^ parasites exhibited extensive cross-resistance to buparvaquone, a structural analog of atovaquone (Hudson et al., 1985), in both *T. gondii* (∼223-fold; Fig. 7b) and *P. falciparum* (∼106-fold; Fig. 8c). Notably, we found minimal cross-resistance to the other tested compounds (maximum 4-fold change in resistance to trifloxystrobin; Table 1). For example, ATV^R^ *P. falciparum* parasites have only mild cross-resistance, and ATV^R^ *T. gondii* parasites have slightly increased sensitivity, to the strobilurin compounds trifloxystrobin and azoxystrobin (Fig. 7d-e; Fig. 8e-f; Table 1). Strobilurins have been shown to target the Q_o_ site of Complex III in fungi (Bartlett et al., 2002), and a study that introduced *P. falciparum* Q_o_ site residues into the yeast Q_o_ site indicated that azoxystrobin may also target this site in apicomplexans (Vallieres et al., 2013). Given the small shifts in IC_50_ observed, our data suggest that if the strobilurins bind the Q_o_ site, they may do so in a different manner to atovaquone and buparvaquone. The chemically diverse compounds that we identified in our screen may, therefore, be useful in the treatment of ATV^R^ parasitic infections. However, we note that several other Q_o_ site mutations can confer atovaquone resistance (Korsinczky et al., 2000, McFadden et al., 2000, Srivastava et al., 1999), and as such further studies could test whether these compounds are effective against other ATV^R^ strains.

Our screen identified two compounds that, to our knowledge, have not been characterized as ETC inhibitors before. The first of these is MMV024397 (6-(4-Benzylpiperidin-1-yl)-*N*-cyclopropylpyridine-3-carboxamide), a compound that is listed under the ‘malaria’ disease set of the MMV ‘Pathogen Box’ and shown to inhibit the proliferation of *P. falciparum* (Fig. 2g) (Tougan et al., 2019), but for which very little other information exists. We demonstrated that MMV024397 inhibited ETC function in both *T. gondii* and *P. falciparum* in a manner consistent with Complex III inhibition. Future studies exploring exactly how this compound inhibits Complex III are warranted.

The second novel ETC inhibiting compound we identified is the aminopyrazole carboxamide compound MMV688853, which has been characterized previously as an inhibitor of *Tg*CDPK1 (Zhang et al., 2014, Huang et al., 2015). The Huang *et al.* (2015) study generated a parasite strain in which the ‘gatekeeper’ residue of *Tg*CDPK1 was mutated (*Tg*CDPK1^G128M^) to render *Tg*CDPK1 resistant to aminopyrazole carboxamides. They found that, despite this mutation, parasite proliferation could still be impaired by several aminopyrazole carboxamide derivatives of MMV688853, suggesting a second target. Our data reveal that the second target of MMV688853 is Complex III of the ETC. Given that we observe no noticeable shift in the IC_50_ of MMV688853 in parasites where *Tg*CDPK has been engineered to be resistant to this compound (Fig. 4f), our data suggest that Complex III is a major target of MMV688853 in the parasite. Mutations in cytochrome *b* can lead to the rapid emergence of resistance to Complex III inhibitors such as atovaquone (McFadden et al., 2000), and it will be of interest to explore whether the dual-targeting properties of MMV688853 make *T. gondii* less prone to developing resistance. It will also be of interest to screen other aminopyrazole carboxamide compounds and/or perform structure-activity relationship studies to determine the chemical basis for MMV688853’s dual inhibition of *Tg*CDPK and Complex III.

We found that MMV688853 failed to inhibit the proliferation (Fig. 2h) and oxygen consumption (Fig. 6g; Fig. S4f) of *P. falciparum* at the concentration ranges we tested (up to 40 μM for proliferation and 50 μM for oxygen consumption). The difference in activity of this compound against *T. gondii* and *P. falciparum* is curious. It is conceivable that these differences are due to impaired uptake of MMV688853 into *P. falciparum* parasites. However, given that we performed the OCR assays with plasma membrane-permeabilized *P. falciparum* parasites (Fig. 6g; Fig. S4f), this explanation is unlikely. A previous study found that MMV688853 was particularly potent against the ookinete stage of *P. berghei* (IC_50_ 220 nM) (Calit et al., 2018). The ookinete is the motile zygote that forms in the midgut of the mosquito vector shortly after transmission of the parasite from the vertebrate host. The potency of MMV688853 against ookinetes was suggested to result from its targeting the *Plasmodium* homolog of *Tg*CDPK1, which is proposed to play a key role in transmission of the parasite into the insect stages of the life cycle (Billker et al., 2004). However, given its dual activity, it is also conceivable that MMV688853 targets the ETC of *Plasmodium*, which becomes more important in the insect stages of the parasite life cycle (Ke et al., 2019, Hino et al., 2012). At odds with this hypothesis is that Complex III is essential in both insect and vertebrate life stages of *Plasmodium* (Ke et al., 2019, Hino et al., 2012, Painter et al., 2007). A final possibility is that MMV688853 targets a site on *T. gondii* Complex III that is not conserved (or potentially not accessible) in Complex III in *P. falciparum* parasites. Whether there are structural differences between Complex III in *T. gondii* and *P. falciparum* that could explain the insensitivity of *P. falciparum* parasites to MMV688853 remains to be seen, but will be a priority for future research.

In summary, our work has developed a scalable pipeline to screen compound libraries to identify inhibitors of the ETC in apicomplexan parasites and characterize their targets. We identified chemically diverse Complex III inhibitors, including MMV688853, which our data suggest is a dual Complex III and *Tg*CDPK1 inhibitor. As many of the identified Complex III inhibitors were active against atovaquone-resistant *T. gondii* and *P. falciparum*, these findings will aid in the development of much-needed new therapeutics against these parasites.

## Materials and methods

### Host cell and parasite culture, and genetic manipulation

Tachyzoite-stage *T. gondii* parasites were cultured in human foreskin fibroblasts (HFF) in Dulbecco’s modified Eagle’s medium (DMEM) containing 2 g/L NaHCO_3_, supplemented with 1% (v/v) fetal calf serum, 50 units/mL penicillin, 50 μg/mL streptomycin, 10 μg/mL gentamicin, 0.25 μg/mL amphotericin B, and 0.2 mM L-glutamine. RH strain *T. gondii* parasites expressing the tandem dimeric Tomato (tdTomato) red fluorescent protein (Chtanova et al., 2008) were used in the initial drug screening assays and for most subsequent *T. gondii* experiments. For the atovaquone resistance experiments, we used wild type ME49 strain parasites or atovaquone-resistant ME49 strain parasites (clone R32), both described previously ((McFadden et al., 2000); a kind gift from Michael Panas and John Boothroyd, Stanford University). To allow us to undertake fluorescence proliferation assays with these ME49 strain parasites, we introduced a tdTomato-encoding vector (Rajendran et al., 2017) into these lines.

To introduce a glycine to methionine mutation at residue 128 of the *Tg*CDPK1 protein of *T. gondii* parasites (*Tg*CDPK1^G128M^), we used a CRISPR-Cas9-based genome editing strategy. We introduced a single guide RNA (sgRNA) targeting the desired region of the open reading frame of the *tg*cdpk1 gene into the pSAG1::Cas9-U6-UPRT vector (Addgene plasmid 54467; (Shen et al., 2014)) using Q5-site directed mutagenesis according to the manufacturer’s instructions (New England Biolabs). We performed the Q5 reaction using the following primers 5´-AAAGGCTACTTCTACCTCGTGTTTTAGAGCTAGAAATAGCAAG-3´ and 5´-AACTTGACATCCCCATTTAC-3´. We also generated a double stranded donor DNA encoding the *Tg*CDPK1^G128M^ mutation flanked by 42-45 bp of homologous flanks to either side of the target site. To do this, we annealed the oligonucleotides 5´-CTGTATGAATTCTTCGAGGACAAAGGCTACTTCTACCTCGTCatgGAAGTGTACAC GGGAGGCGAGTTGTTCGACGAGATCATTTCCCGC-3´ and 5´-GCGGGAAATGATCTCGTCGAACAACTCGCCTCCCGTGTACACTTCcatGACGAGGTAGAAGTAGCCTTTGTCCTCGAAGAATTCATACAG-3´ (mutated base pairs are indicated by the lower case letters). We combined the sgRNA expressing plasmid (which also encodes Cas9-GFP) and donor DNA and transfected them into TATiΔ*ku80*/Tomato^+^ parasites by electroporation as described previously (Jacot, 2020). Two days after transfection, we selected and cloned GFP^+^ parasites by flow cytometry. We PCR-amplified the genomic DNA of several clones using the primers 5´-AGTGAAGCAGAAGACGGACAAG-3´ and 5´-GAGGTCCCGATGTACGATTTTA-3´, and checked for successful modification by Sanger sequencing. We termed the resulting parasite strain ‘*Tg*CDPK1^G128M^’.

3D7 strain *P. falciparum* parasites were maintained in synchronous continuous culture in Roswell Park Memorial Institute (RPMI)-1640 medium supplemented with 25 mM HEPES, 20 mM D-glucose, 200 μM hypoxanthine, 24 mg/L gentamicin and Albumax II (0.6% w/v), as described previously (de Villiers et al., 2013, Allen and Kirk, 2010). Atovaquone-resistant parasites were generated by maintaining cultures at 1% parasitaemia in the presence of atovaquone at an initial concentration equivalent to the IC_50_ of atovaquone (0.5 nM). Fresh medium, erythrocytes and atovaquone were added every 2 days and parasitaemia was adjusted to 1%. The atovaquone concentration was increased by 0.5 nM every week for 12 weeks. Once parasites were proliferating in the presence of 10 nM atovaquone (∼20× IC_50_), clonal populations were selected by limiting dilution cloning. We PCR-amplified the cytochrome *b* gene of *P. falciparum* using primers described previously (Goodman et al., 2016): 5´-CTCTATTAATTTAGTTAAAGCACAC-3´ and 5´-ACAGAATAATCTCTAGCACC-3´. We checked for mutations in the amplified cytochrome *b* gene by Sanger sequencing using the following primers: 5´-AGCAGTAATTTGGATATGTGGAGG-3´ and 5´-AATTTTTAATGCTGTATCATACCCT-3´. 3D7 strain *P. falciparum* parasites expressing yeast dihydroorotate dehydrogenase (yDHODH) were a kind gift from Emily Crisafulli and Stuart Ralph (University of Melbourne), and were maintained on 10 nM WR99210 (which was removed prior to growth assays) as described previously (Dickerman et al., 2016).

### Compounds

The ‘Pathogen Box’ compounds were kindly provided by MMV in 96-well plates containing 10 mM stock solutions dissolved in DMSO. Additional amounts of several compounds were purchased from Sigma Aldrich and dissolved in DMSO (stock concentration given in brackets), including azoxystrobin (31697-100MG; 50 mM), trifloxystrobin (46447-100MG; 50 mM), auranofin (A6733-10MG; 50 mM), buparvaquone (SML1662-25MG; 3 mM), and atovaquone (A7986-10MG; 10 mM). 3MB-PP1 was purchased from Cayman Chemical (17860; 10 mM). Additional MMV688853 (BKI-1517; 10 mM) was a kind gift from Wes van Voorhis (University of Washington). Additional MMV024397 was also provided by MMV. The DMSO concentration introduced when using these compounds in assays was < 0.2% (v/v), except MMV688853 when used at the higher concentrations (up to 50 μM) in the *Plasmodium* assays (up to 0.5% (v/v) DMSO).

### Screening compounds using Seahorse XFe96 extracellular flux assay

The MMV ‘Pathogen Box’ compounds were screened for their ability to inhibit O_2_ consumption of intact *T. gondii* parasites using a Seahorse XFe96 flux assay described previously (Hayward et al., 2022) with slight modifications. Parasites (tdTomato-expressing RH strain *T. gondii* tachyzoites) were mechanically egressed from host cells by passing them through a 26-gauge needle, then filtered through a 3 µm polycarbonate filter to remove host cell debris, counted using a hemocytometer, and pelleted by centrifugation (1500 × *g*, 10 min, RT). The medium was aspirated and parasites were washed once in Base Medium (Agilent) supplemented with 1 mM L-glutamine and 5 mM D-glucose (termed supplemented Base Medium), then resuspended in supplemented Base Medium to 1.5 × 10^7^ parasites/mL. Parasites (1.5 × 10^6^) were seeded into wells of a Seahorse XFe96 cell culture plate coated with 3.5 μg/cm^2^ CellTak cell adhesive (Corning) and attached to the bottom by centrifugation (800 × *g* for 3 min). The final well volume was 175 μL, achieved by adding supplemented Base Medium. MMV ‘Pathogen Box’ compounds were prepared such that the final concentration upon injection (25 μL injection volumes) would be 1 μM (8 μM for compounds to be injected from port A; 9 μM for compounds to be injected from port B; and 10 μM for compounds to be injected from port C). During the XFe96 assay, three compounds were sequentially injected into each well (from ports A-C) and the OCR measured for three cycles of 30 s mixing followed by 3 min measuring. A final injection of the known ETC Complex III inhibitors antimycin A (10 μM) and atovaquone (1 μM) from port D was used as a control to validate that the assay was measuring mitochondrial OCR, and to enable determination of non-mitochondrial OCR. In instances where ‘hit’ compounds were injected from ports A or B, compounds injected from later ports in that particular well were retested in a subsequent assay to ensure compounds injected after the ‘hit’ compound were not missed. Percent inhibition of OCR by each of the 400 compounds was calculated relative to the antimycin A- and atovaquone-treated control (set to 100% inhibition). An arbitrary cut-off of >30% inhibition of OCR was applied in selecting candidate ETC inhibitors from the screen.

### Seahorse XFe96 extracellular flux analysis of intact *T. gondii* parasites

The inhibitory activity of selected MMV ‘Pathogen Box’ compounds against the OCR of intact *T. gondii* parasites was assessed using a previously described Seahorse XFe96 flux assay (Hayward et al., 2022) with slight modifications. *T. gondii* parasites were prepared and seeded into wells of a Cell-Tak coated Seahorse XFe96 cell culture plate as described above. The final well volume was 175 μL, achieved with supplemented Base Medium. Carbonyl cyanide 4-(trifluoromethoxy)phenylhydrazone (FCCP) was prepared in Base Medium such that the final concentration upon injection would be 1 μM (8 μM for injection from port A). A serial dilution of the test compounds as well as a no-drug (DMSO) control was performed in supplemented Base Medium, and loaded into port B at 9× the desired final concentrations. Supplemented Base Medium was injected from port C, and a final injection of the known ETC Complex III inhibitor atovaquone (5 μM final concentration) from port D was used as a control to completely inhibit mitochondrial OCR. The OCR and ECAR were measured for three cycles of 30 s mixing followed by 3 min measuring at baseline after injections from port A and port C, and for six cycles of 30 s mixing followed by 3 min measuring after injections from port B and port D. Mitochondrial OCR (mOCR) was calculated by subtracting the last OCR reading after atovaquone injection (port D) from the last OCR reading after test compound injection (port B). Percent mOCR relative to the drug-free control was plotted against the test compound concentration, and a sigmoidal four parameter logistic (4PL) curve was fitted using nonlinear regression in GraphPad Prism to yield the compound concentration required for 50% inhibition (IC_50_) of *T. gondii* OCR.

### Seahorse XFe96 extracellular flux analysis of plasma membrane-permeabilized parasites

Measurement of substrate-elicited OCR of digitonin-permeabilized *T. gondii* parasites was performed as described previously (Hayward et al., 2021, Hayward et al., 2022). Briefly, freshly egressed *T. gondii* parasites were passed through a 3 μm filter to remove host cell debris, counted using a hemocytometer, and pelleted by centrifugation (1500 × *g*, 10 min, RT). Parasites were washed once in non-supplemented Base Medium, resuspended in non-supplemented Base Medium to 1.5 × 10^7^ parasites/mL and incubated at 37°C for approximately 1 hour to deplete endogenous substrates. Parasites (1.5 × 10^6^) were added to the wells of a Cell-Tak-coated Seahorse cell culture plate and centrifuged (800 × *g*, 10 min, RT) to adhere parasites to the bottom of the wells. Just before the beginning of the assay, Base Medium was removed and replaced with 175 μL mitochondrial assay solution (MAS) buffer (220 mM mannitol, 70 mM sucrose, 10 mM KH_2_PO_4_, 5 mM MgCl_2_, 0.2% (w/v) fatty acid-free bovine serum albumin (BSA), 1 mM EGTA and 2 mM HEPES-KOH pH 7.4) containing 0.002% (w/v) digitonin to permeabilize the parasite plasma membrane. The following compounds were prepared in MAS buffer (final concentration after injection given in brackets) and loaded into ports A-D of the XFe96 sensor cartridge: Port A, ETC substrates malate (Mal; 10 mM) or sn-glycerol 3-phosphate bis(cyclohexylammonium) salt (G3P; 25 mM) plus FCCP (1 μM); Port B, the test compounds atovaquone (1.25 μM), auranofin (10 μM), azoxystrobin (20 μM), trifloxystrobin (2.5 μM), MMV688853 (20 μM), buparvaquone (5 μM) or MMV024397 (20 μM); Port C, *N*,*N*,*N*´,*N*´-tetramethyl-*p*-phenylenediamine dihydrochloride (TMPD; 0.2 mM) mixed with ascorbic acid (3.3 mM); Port D, sodium azide (NaN_3_; 10 mM). The OCR was assessed for three cycles of 30 s mixing followed by 3 min measuring to establish baseline OCR before substrate injection, for three cycles of 30 s mixing followed by 3 min measuring after the injection of substrates from ports A and C, and for six cycles of 30 s mixing followed by 3 min measuring after the injections of compounds from ports B and D. A minimum of four background wells (containing no parasites) were used in each plate, and 3 technical replicates were used for each condition.

OCR measurements of digitonin-permeabilized *P. falciparum* parasites were performed using a protocol modified from one described previously (Sakata-Kato and Wirth, 2016). On the day of the assay, 200 mL of *P. falciparum* culture at 4% (v/v) hematocrit and at least 5% parasitaemia was enriched for trophozoites by passing through a MACS CS column placed in the magnetic field of a SuperMACS II (Miltenyi Biotec) separator according to the manufacturer’s instructions. The trophozoites were freed from erythrocytes by treating with 0.05% (w/v) saponin at 37 °C for 5 minutes. The obtained parasite pellets were washed with phosphate buffered saline (PBS) until the supernatant was no longer red (*i.e.* until most host cell hemoglobin had been removed). Parasites were counted using a hemocytometer and prepared at 5×10^7^ parasites/mL in MAS buffer supplemented with 10 mM malate and 0.002% (w/v) digitonin. Parasites were seeded at a density of 5×10^6^ cells in a Cell-Tak-coated XFe96 cell culture plate and centrifuged (800 × *g*, 10 min, RT) to adhere the parasites to the bottom of the wells. Supplemented MAS buffer (75 μL) was carefully added to the wells without disturbing the cell monolayer. The following compounds were prepared in MAS buffer (final concentration after injection given in brackets) and loaded into ports A-C of the XFe96 sensor cartridge: Port A, a 2-fold serial dilution of the test compounds; Port B, TMPD (0.2 mM) mixed with ascorbic acid (2 mM); Port C, sodium azide (NaN_3_; 10 mM). The OCR was measured for five cycles of 20 s mixing, 1 min waiting, 2.5 min measuring at baseline and after each injection. Percent mOCR relative to the drug-free control was plotted against the test compound concentration, and a variable slope (four parameters) curve was fitted using nonlinear regression in GraphPad Prism to yield the IC_50_ for *P. falciparum* OCR.

### *T. gondii* fluorescence proliferation assays

The anti-parasitic activity of selected MMV ‘Pathogen Box’ compounds was assessed by fluorescence proliferation assays, measuring the proliferation of tdTomato-expressing *T. gondii* parasites as described previously (Rajendran et al., 2017). Briefly, 2000 parasites were added to wells of a clear bottom, black 96-well plate containing HFF cells, in phenol red-free DMEM supplemented with 1% (v/v) fetal calf serum, 50 units/mL penicillin, 50 μg/mL streptomycin, 10 μg/mL gentamicin, 0.25 μg/mL amphotericin B, and 0.2 mM L-glutamine. A serial dilution of the desired compounds was performed and added to wells of the plate. Parasites were allowed to proliferate and fluorescence was measured daily using a FLUOstar OPTIMA Microplate Reader (BMG LABTECH). Percent parasite proliferation relative to the no-drug control at mid-log phase was plotted against the compound concentration, and a variable slope (four parameters) curve was fitted using nonlinear regression in GraphPad Prism, enabling calculation of the IC_50_ of compound against *T. gondii* proliferation.

### *P. falciparum* proliferation assays

The anti-plasmodial activity of selected MMV ‘Pathogen Box’ compounds was assessed using a SYBR Safe-based fluorescence assay described previously (Smilkstein et al., 2004, Spry et al., 2013). Assays were set up using ring-stage *P. falciparum*-infected erythrocytes in culture medium at a hematocrit of 1% and parasitemia of 0.5%. Parasites were allowed to proliferate for 96 h, after which the percentage parasite proliferation was plotted against the compound concentration. A variable slope (four parameters) curve was fitted to the data using nonlinear regression in GraphPad Prism, enabling calculation of the IC_50_ of the compound against *P. falciparum* proliferation.

### Flow cytometry analysis of *T. gondii* viability

Freshly egressed RHΔ*hxgprt* strain *T. gondii* parasites were passed through a 3 μm filter to remove host cell debris. Parasites were pelleted by centrifugation (1500 × *g*, 10 min, RT) and resuspended in phenol red-free DMEM containing 5 mM D-glucose and 1 mM L-glutamine. Parasites were incubated (37°C, 5% CO_2_) for various times (15 to 120 min) in the presence of DMSO (vehicle control), auranofin (1 μM, 20 μM or 100 μM) or atovaquone (10 μM). Propidium iodide (PI, 15 μM) was then added and parasites were incubated for a further 20 min (RT, protected from light), before being analyzed on a BD LSR II Flow Cytometer. FSC and SSC parameters were used to gate for single parasites. PI fluorescence was excited using the 488 nm laser and detected with a 670/14nm filter. Acquired data were exported for further analysis using FlowJO 10 (BD) software.

### Complex III enzymatic assay

To measure Complex III enzymatic activity in *T. gondii*, we adapted an assay previously established for mammalian cells (Spinazzi et al., 2012). Egressed parasites were passed through a 5 µm polycarbonate filter to remove host cell debris, counted using a hemocytometer, and pelleted by centrifugation (10 min, 1500 × *g,* RT). Pellets were washed in 1 mL cold PBS and centrifuged (1 min, 12000 × *g,* RT). Parasites were resuspended to 2.5 × 10^8^ parasites/mL in MAS buffer containing 0.2% (w/v) digitonin, and lysed on a spinning wheel (30 min, 4°C). Complex III assay buffer (25 mM KH_2_PO_4_ pH 7.5, 75 μM oxidised equine heart cytochrome *c*, 100 μM EDTA, 0.025% (v/v) Tween-20 and 1.21 mM sodium azide) was prepared and aliquoted into the wells of a 24-well plate. The following compounds (or DMSO as a no-drug vehicle control) were added to three wells each at the indicated final concentrations: atovaquone (1.25 μM), auranofin (10 μM), azoxystrobin (20 μM), trifloxystrobin (2.5 μM), MMV688853 (20 μM), buparvaquone (5 μM), MMV024397 (20 μM).

A baseline reading was taken by measuring the absorbance at 550 nm every 15 s for 2 min using a TECAN Infinite 200 PRO plate reader warmed to 37°C. Parasite lysate (an equivalent of 6.25 × 10^6^ parasites per mL) was then added to two of the three wells per drug (duplicate technical experimental wells) while MAS buffer was added to the remaining well (as a ‘no parasite lysate’ background control), and a further baseline reading was taken every 15 s for 2 min. To start the reaction, 5 μM reduced decylubiquinol in DMSO was added to each well, and absorbance at 550 nm was measured every 15 s for 60 min.

To calculate enzymatic activity, absorbance was plotted as a function of time. The initial rate was estimated from the first 5 minutes after adding decylubiquinol, and divided by the extinction coefficient for reduced cytochrome *c* (18.5 mM^−1^ cm^−1^) according to the Beer-Lambert Law. For each condition, the background (initial rate in the absence of parasite lysate) was subtracted from the observed value to yield the calculated activity.

### *T. gondii* invasion assay

To determine the effects of compounds on parasite invasion, we undertook invasion assays based on a modified version of a previously described protocol (Kafsack et al., 2004). TATiΔ*ku80*/Tomato^+^ strain *T. gondii* parasites were cultured in HFF cells such that most parasites were still intracellular prior to the assay. Extracellular parasites were removed by washing the flask three times with warm intracellular (IC) buffer (5 mM NaCl, 142 mM KCl, 2 mM EGTA, 1 mM MgCl_2_, 5.6 mM D-glucose and 25 mM HEPES, pH 7.4). Infected host cells were then scraped from the flasks, passed through a 26-gauge needle to mechanically egress the parasites, and filtered through a 3 μm polycarbonate filter to remove host cell debris. Parasites were counted using a hemocytometer and diluted to 5 × 10^5^ parasites per mL in IC buffer with either DMSO (vehicle control), MMV688853 (5 μM), 3MB-PP1 (5 μM) or atovaquone (1 μM), added to wells of a 24-well plate containing confluent HFF cells cultured on coverslips, and incubated at 37°C for 45 min to allow parasites to attach to host cells. To induce invasion, IC buffer was removed and replaced with DMEM containing DMSO/drug added at the above concentrations. Parasites were allowed to invade for 25 min at 37°C, before being fixed in 3% (w/v) paraformaldehyde (PFA) and 0.1% (w/v) glutaraldehyde in PBS for 20 min at RT. After fixation, coverslips were blocked in 2% (w/v) BSA in PBS. To identify uninvaded extracellular parasites, we conducted immunofluorescence assays. We labelled uninvaded extracellular parasites with the *T. gondii* cell surface marker mouse anti-SAG1 primary antibody (Abcam, Ab8313; 1:1000 dilution) and a goat anti-mouse Alexa Fluor 488 Plus secondary antibody (Thermo Fisher Scientific, A32723; 1:500 dilution). Coverslips were mounted onto slides, the identity of the samples blinded to the observer, and invaded vs non-invaded parasites were quantified on a DeltaVision Elite deconvolution microscope (GE Healthcare) fitted with a 100× UPlanSApo oil immersion objective lens (NA 1.40). Parasites that were both red (Tomato^+^) and green (SAG1^+^) were considered to be extracellular, while those that were red but not green were considered as having invaded a host cell. At least 100 parasites were counted per condition.

### *T. gondii* intracellular proliferation assay

TATiΔ*ku80*/Tomato^+^ strain *T. gondii* parasites were prepared in a similar way to the invasion assay. Following mechanical egress in IC buffer, parasites were counted and diluted to 5 × 10^4^ parasites/mL in IC buffer, added to wells of a 24-well plate containing confluent HFF cells cultured on coverslips, and incubated at 37°C for 45 min to allow the parasites to attach to host cells. IC buffer was removed and replaced with 1 mL DMEM, and parasites were allowed to invade and begin to proliferate for 4 h at 37°C in the absence of drug. Medium was then removed, cells were washed twice to remove uninvaded parasites, and replaced with 1 mL DMEM with either DMSO (vehicle control), MMV688853 (5 μM), 3MB-PP1 (5 μM) or atovaquone (1 μM). Parasites were cultured for a further 19 h at 37°C, then fixed in 3% (w/v) PFA in PBS for 15 min. Coverslips were mounted onto slides, and the identity of each was blinded to the observer. The number of parasites per vacuole were quantified on a DeltaVision Elite deconvolution microscope (GE Healthcare). At least 100 vacuoles were counted per condition.

## Acknowledgements

This work was supported by a Research School of Biology innovation grant to ER, DC, AGM and GGvD, a National Health and Medical Research Council Ideas Grant (GNT1182369) to GGvD, AGM and KJS, an Australian Research Council Discovery project (DP1801032) to AGM, and an Australian Government Research Training Program Scholarship to JAH. The funders had no role in study design, data collection and analysis, decision to publish, or preparation of the manuscript. The authors declare that they have no conflicts of interest.

We would like to thank the Medicines for Malaria Venture for supplying the ‘Pathogen Box’ compounds, Wes van Voorhis (University of Washington) for supplying extra MMV688853 (BKI-1517) compound, John Boothroyd and Michael Panas (Stanford University) for supplying atovaquone-resistant *T. gondii* parasites, Emily Crisafulli and Stuart Ralph (University of Melbourne) for supplying yDHODH-expressing *P. falciparum* parasites, Harpreet Vohra and Michael Devoy (ANU) for assistance with flow cytometry, Michael Devoy for assistance with establishing the XFe96 assays, Teresa Neeman from the ANU Statistical Consulting Unit for assistance with data analysis, the 2020 ANU Parasitology Course (BIOL3142) for contributing to trial experiments with MMV688853, Adele Lehane (ANU) and the ANU parasitology journal club for comments on the manuscript, and the Canberra Branch of the Australian Red Cross Lifeblood for the provision of erythrocytes.

## Supplementary Figures

**Supplementary Figure 1.**
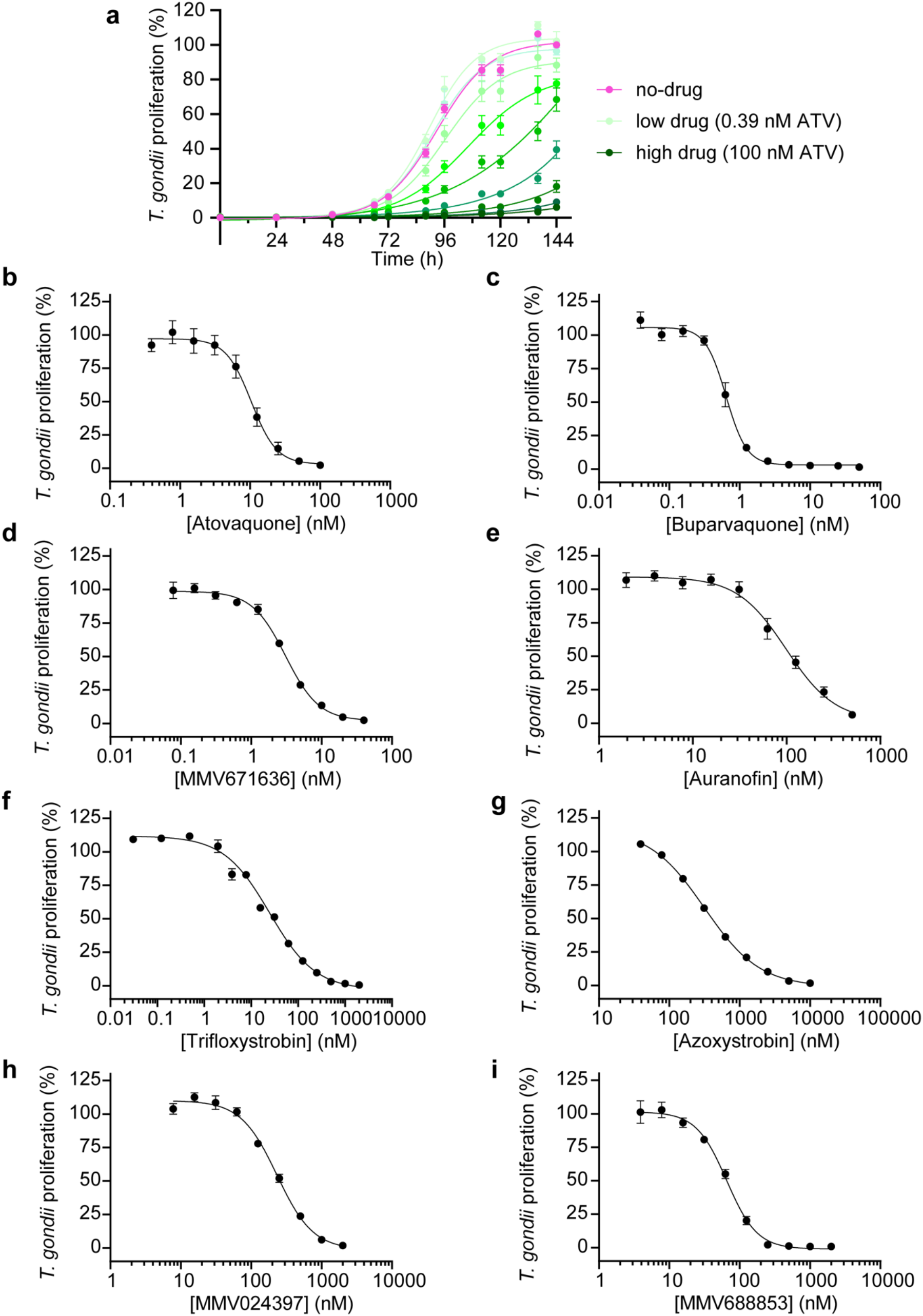
Candidate ETC inhibitors inhibit proliferation of *T. gondii* parasites. **(a)** Proliferation of tdTomato-expressing *T. gondii* parasites cultured in the absence of drug (pink circles), or in the presence of atovaquone (two fold serial dilution from highest concentration (100 nM; dark green) to lowest concentration (0.39 nM; light green)) over a 6-day period. Values are expressed as a percent of the average fluorescence from the no-drug control on the final day of the experiment, and represent the mean ± SD of three technical replicates. Data are from one experiment and are representative of three independent experiments. Similar proliferation curves were obtained for each test compound but are not shown. **(b-i).** Dose-response curves depicting the percent of *T. gondii* parasite proliferation in the presence of a range of concentrations of **(b)** atovaquone, **(c)** buparvaquone, **(d)** MMV671636, **(e)** auranofin, **(f)** trifloxystrobin, **(g)** azoxystrobin, **(h)** MMV024397, or **(i)** MMV688853. Values are expressed as a percent of the average fluorescence from the no drug control at mid-log phase growth, and represent the mean ± SEM of three independent experiments, each conducted in triplicate; error bars that are not visible are smaller than the symbol.

**Supplementary Figure 2.**
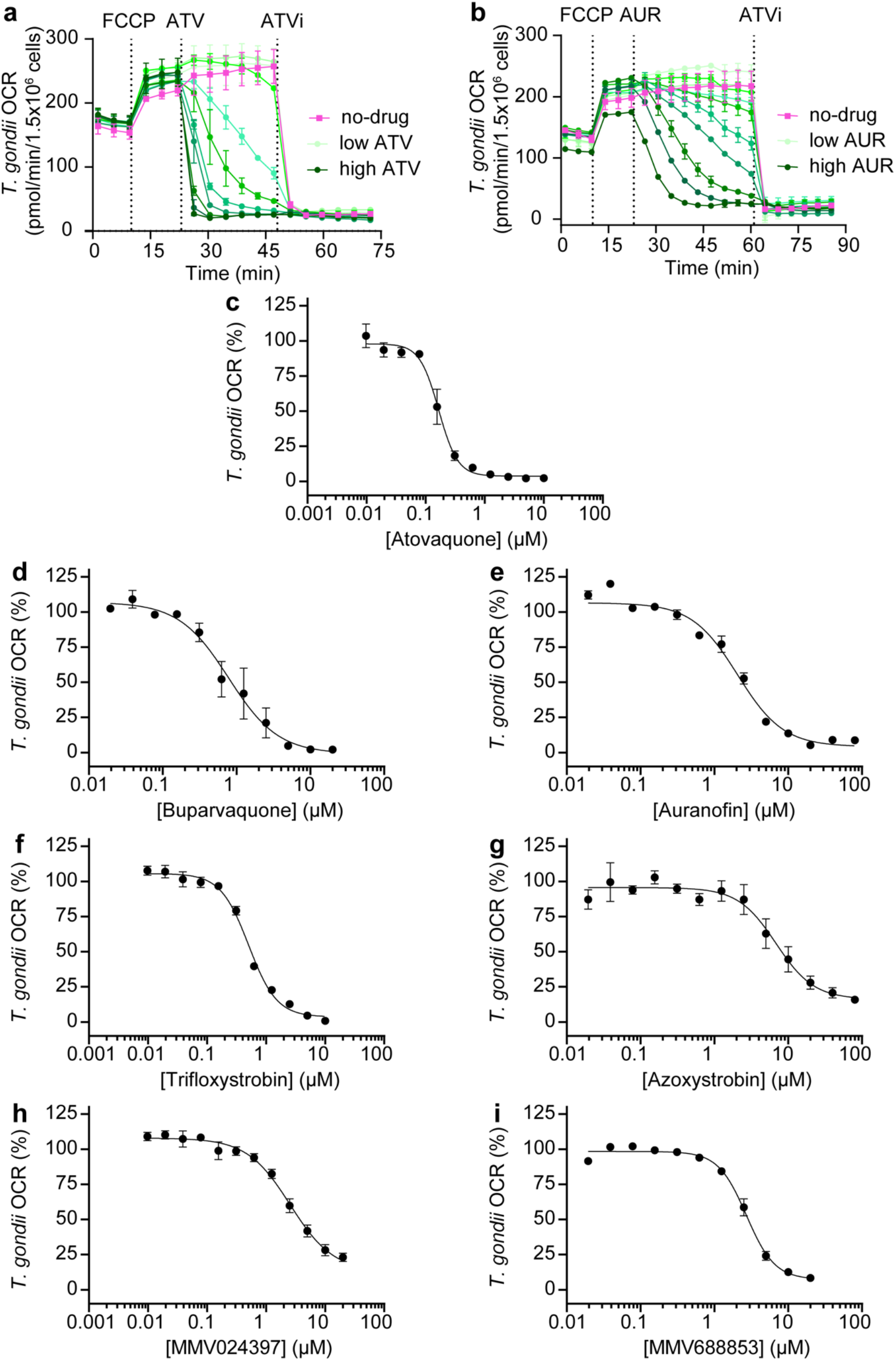
Identified compounds inhibit O*_2_* consumption in *T. gondii*. **(a-b)** Traces depicting the changes in oxygen consumption rate (OCR) over time of intact *T. gondii* parasites incubated with no drug (pink) or with **(a)** atovaquone (ATV) (two fold serial dilution from highest concentration (10 μM; dark green) to lowest concentration (0.01 μM; light green)) or **(b)** auranofin (AUR) (two fold serial dilution from highest concentration (80 μM; dark green) to lowest concentration (0.08 μM; light green)). FCCP (1 μM) was injected into the well to uncouple electron transport from ATP synthesis and thus elicit the maximal OCR. A range of concentrations of the test compounds were then injected and the inhibition of OCR measured over time. A final injection of an inhibitory concentration of atovaquone (ATVi; 5 µM) fully inhibited mitochondrial OCR. Values represent the mean ± SD of two technical replicates from a single experiment and are representative of three independent experiments. Similar OCR inhibition traces were obtained for each test compound but are not shown. **(c-i)** Dose-response curves depicting the percent of *T. gondii* OCR in the presence of increasing concentrations of **(c)** atovaquone, **(d)** buparvaquone, **(e)** auranofin, **(f)** trifloxystrobin, **(g)** azoxystrobin, **(h)** MMV024397 or **(i)** MMV688853. Values represent the percent OCR relative to the no-drug (100% OCR) and inhibitory atovaquone-treated (0% OCR) controls, and depict the mean ± SEM of three independent experiments, each conducted in duplicate; error bars that are not visible are smaller than the symbol.

**Supplementary Figure 3.**
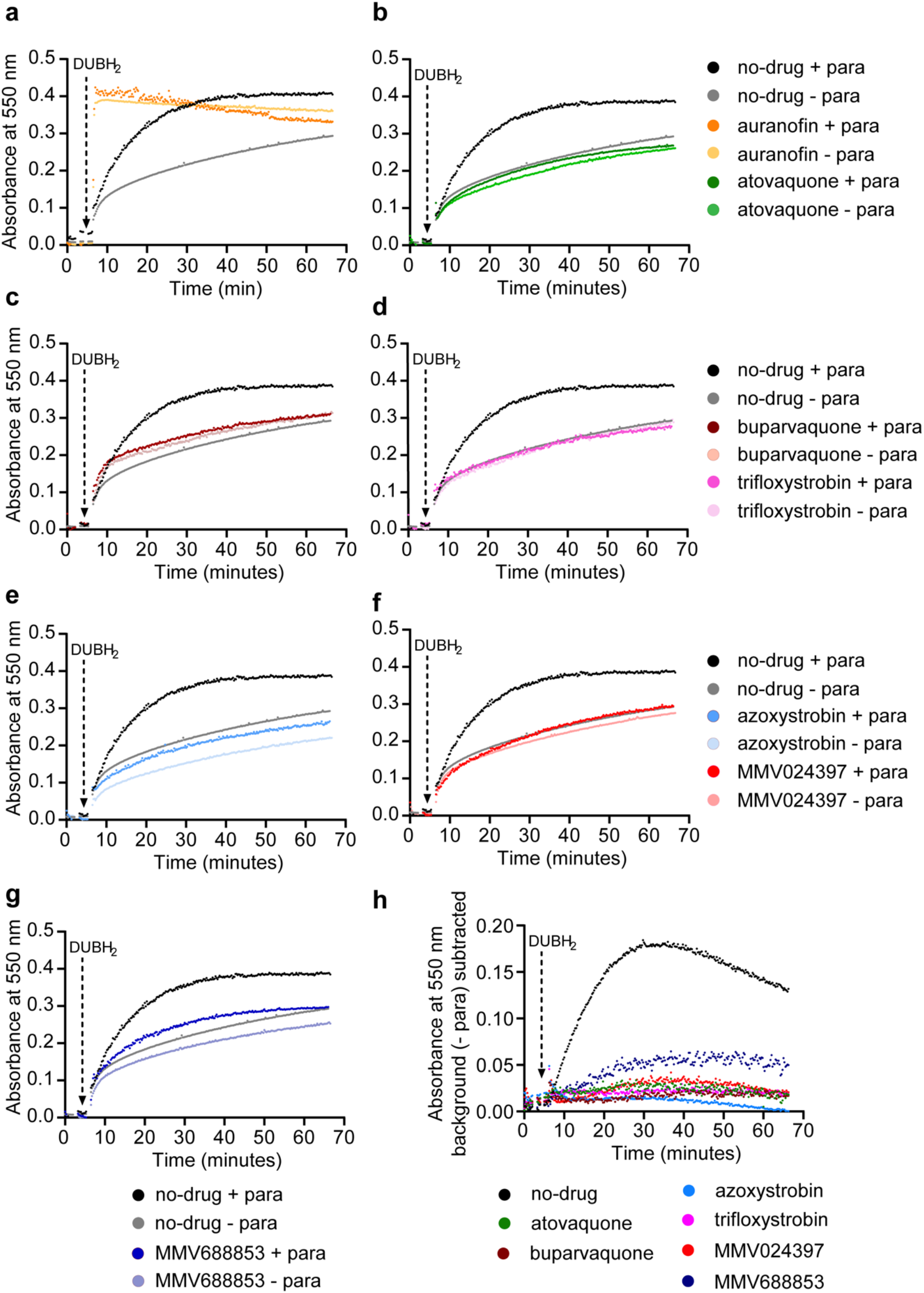
Characterizing Complex III inhibition by the candidate ETC inhibitors in *T. gondii*. **(a-g)** Complex III activity assays showing the change in absorbance of equine heart CytC at 550 nm over time (measured every 15 s) in the presence (+ para, dark shade) or absence (-para, light shade) of parasite extracts, and in the presence of no drug (DMSO vehicle control, black or gray), or **(a)** auranofin (orange, 10 μM), **(b)** atovaquone (green, 1.25 μM), **(c)** buparvaquone (burgundy, 5 μM), **(d)** trifloxystrobin (pink, 2.5 μM), **(e)** azoxystrobin (light blue, 80 μM), **(f)** MMV024397 (red, 20 μM) or **(g)** MMV688853 (dark blue, 20 μM). Data are from a single experiment and are representative of three independent experiments. **(h)** Complex III activity assays showing the change in absorbance of equine heart CytC at 550 nm over time where change in absorbance in the absence of parasite extracts (*i.e.* background absorbance) was subtracted from the change in absorbance in the presence of parasite extracts. Data are from a single experiment and are representative of three independent experiments. Compounds are depicted using the same coloring as in b-g.

**Supplementary Figure 4:**
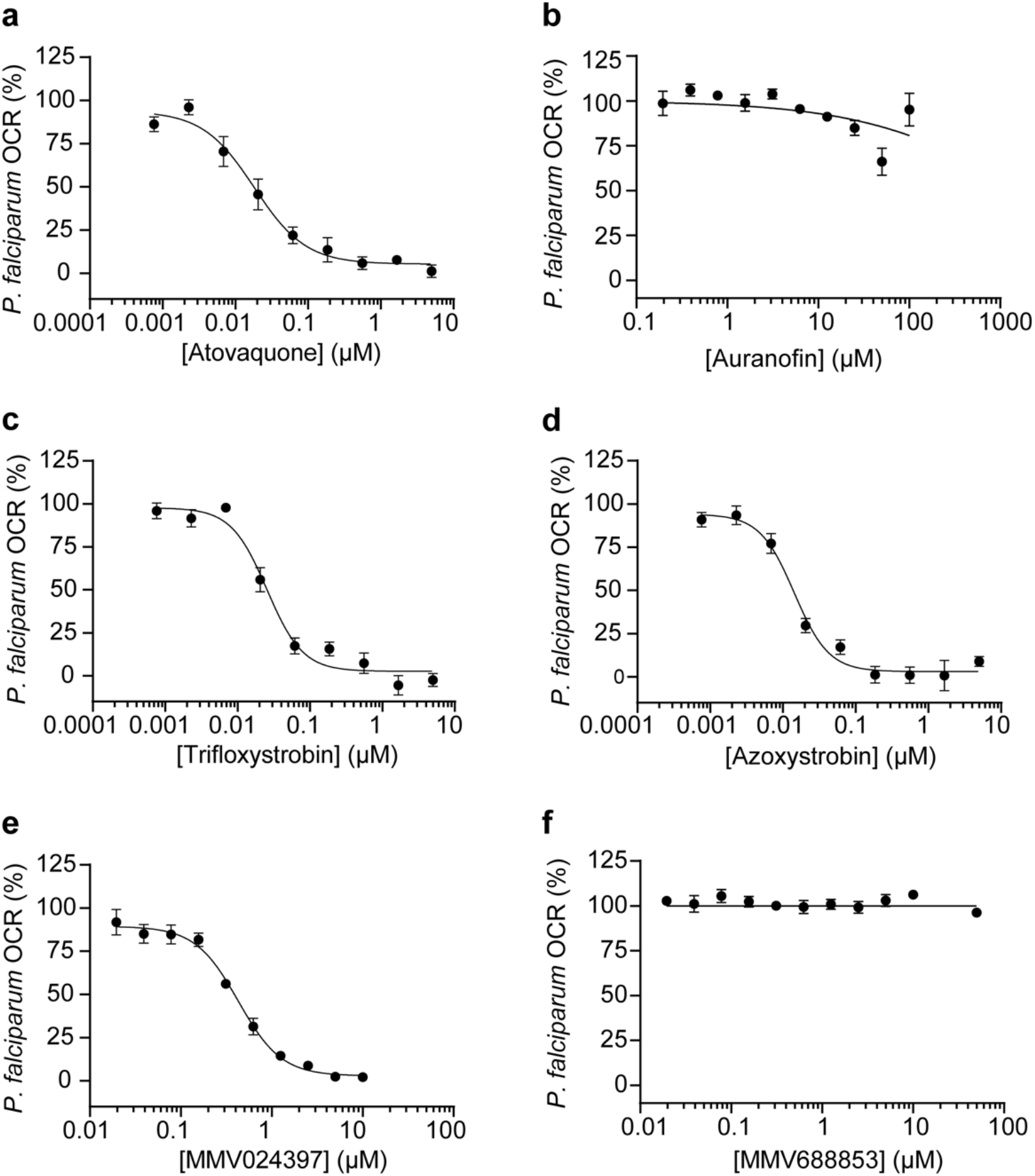
Identified compounds inhibit O*_2_* consumption in *P. falciparum*. **(a-f)** Dose-response curves depicting *P. falciparum* OCR in the presence of increasing concentrations of **(a)** atovaquone, **(b)** auranofin, **(c)** trifloxystrobin, **(d)** azoxystrobin, **(e)** MMV024397 or **(f)** MMV688853. Values represent the percent OCR relative to the no drug (100% OCR) and atovaquone-treated (0% OCR) controls, and are depicted as the mean ± SEM of three independent experiments; error bars that are not visible are smaller than the symbol.

